# Quantification of within-patient *Staphylococcus aureus* phenotypic heterogeneity as a proxy for presence of persisters across clinical presentations

**DOI:** 10.1101/2021.07.27.453929

**Authors:** Julian Bär, Mathilde Boumasmoud, Srikanth Mairpady Shambat, Clément Vulin, Markus Huemer, Tiziano A. Schweizer, Alejandro Gómez-Mejia, Nadia Eberhard, Yvonne Achermann, Patrick O. Zingg, Carlos–A. Mestres, Silvio D. Brugger, Reto A. Schüpbach, Roger D. Kouyos, Barbara Hasse, Annelies S. Zinkernagel

**Affiliations:** Department of Infectious Diseases and Hospital Epidemiology, University Hospital Zurich, University of Zurich, Zurich, Switzerland; Balgrist University Hospital, University of Zurich, Zurich, Switzerland; Clinic for Cardiovascular Surgery, University Hospital Zurich, University of Zurich, Zurich, Switzerland; Institute for Intensive Care Medicine, University Hospital Zurich, University of Zurich, Zurich, Switzerland

## Abstract

**Background:** Difficult-to-treat infections caused by antibiotic susceptible strains have been linked with the occurrence of persisters. Persisters are a subpopulation of dormant bacteria that tolerate antibiotic exposure despite lacking genetic resistance. They can be identified phenotypically upon plating on nutrient agar because of their altered growth dynamics, resulting in colony size heterogeneity. The occurrence of within-patient bacterial phenotypic heterogeneity in various infections and clinical determinants of persister formation remain unknown.

**Methods:** We plated bacteria derived from 132 patient-samples of difficult-to-treat infections directly on nutrient-rich agar and monitored colony growth by time-lapse imaging. Of these, we retained 36 *Staphylococcus aureus* mono-cultures for further analysis. We investigated clinical factors potentially associated with increased colony growth-delay with regression analyses. Additionally, we corroborated the clinical findings using *in vitro* grown static biofilms, exposed to distinct antibiotics.

**Results:** The extent of phenotypic heterogeneity of patient-derived *S. aureus* varied substantially between patients. Increased heterogeneity coincided with increased median growth-delay. Multivariable regression showed that rifampicin treatment was significantly associated with increased median growth-delay. *S. aureus* grown in biofilms and exposed to high concentrations of rifampicin or a combination of rifampicin with either clindamycin or levofloxacin exhibited prolonged growth-delay, correlating with a strain-dependent increase in antibiotic tolerance.

**Conclusions:** Upon direct cultivation on nutrient-rich agar, *S. aureus* from difficult-to-treat infections commonly exhibited colony size heterogeneity. This was due to heterogeneous delays in growth resumption, with delays larger than two days in the most extreme cases. Since bacteria in a dormant state are tolerant to antibiotics, the observation of large growth-delays might have direct clinical implications. Future studies are needed to assess the potential of bacterial phenotypic heterogeneity quantification for staphylococcal infections prognosis.

## Introduction

*Staphylococcus aureus* is frequently part of the normal flora as well as a cause of infections in humans.^1^ Deep-seated infections such as cardiovascular infections and prosthetic joint infections are usually difficult-to-treat, due to the presence of biofilms.^2^ They require prolonged antibiotic treatment and, in some cases, surgical debridement and removal of native or prosthetic material. Despite state-of-the-art treatment, *S. aureus* biofilm-associated infections are linked with increased morbidity and mortality.

Prolonged antibiotic treatment facilitates the emergence of antibiotic resistance, further complicating therapy.^3^ Antibiotic resistance can be preceded by antibiotic tolerance,^4,5^ defined as the ability of an antibiotic susceptible bacterial population to survive a time-limited antibiotic challenge.^6,7^ This property can be conferred either by mutations affecting growth rate or by a phenotypic switch to a dormant state.^8^ The resulting slow- or non-growing bacteria, termed persisters, have been implicated in difficult-to-treat and relapsing *S. aureus* infections.^9–12^

This subpopulation of persisters can be identified upon plating on nutrient-rich agar, because of its altered growth dynamics. In the case of mutations affecting growth rate, stable small colony variants (SCV) can be observed.^13,14^ Yet most often, *S. aureus* isolated from infection sites have been reported to give rise to non-stable small colonies (nsSC), which result from heterogeneous delays in growth resumption of bacterial cells of a clonal population. This heterogeneity in dormancy can be induced by exposure to different stressors such as low pH, antibiotic exposure, biofilm or intracellular environment^15–17,10^ and is reverted when the stress is removed. An infecting strain is likely to encounter most if not all these stressors within a patient, but the mechanisms triggering and regulating the occurrence of persisters in these complex conditions are largely unknown. Few studies monitored colony growth of bacterial populations directly upon recovery from human infection sites.^12,15,18^ We showed that *S. aureus* from abscesses exhibit heterogeneous colony growth-delays *ex vivo*^12,15^ and others reported that *Mycobacterium tuberculosis* colony growth dynamics *ex vivo* change over the course of pulmonary tuberculosis treatment.^18^ These studies focused on single or few clinical cases and do not provide a systematic assessment of phenotypic heterogeneity and persisters at the patient-population level.

Here, we provide the first descriptive epidemiological study quantifying the occurrence of within-patient *S. aureus* phenotypic heterogeneity, as a proxy for presence of persisters, in distinct clinical presentations. We investigated the association of several clinical factors, including antibiotic treatment regimen, with colony growth-delay of *S. aureus* immediately upon isolation from patients. We confirmed the biological effect of specific antibiotics on colony growth-delay using an *in vitro* biofilm assay mimicking the complex environment encountered by bacterial populations *in vivo*.

## Material & Methods

### Study setting and study participants

This study took place between October 2016 and May 2020 at two tertiary care hospitals in Switzerland, the University Hospital of Zurich and the Balgrist University Hospital. Patients with vascular graft /endovascular infections or infective endocarditis were prospectively included in the Vascular Graft Cohort study (VASGRA; KEK-2012-0583) and the Endovascular and Cardiac Valve Infection Registry (ENVALVE; BASEC 2017-01140), respectively. Patients with bone and prosthetic joint infections were included in the framework of the Prosthetic Joint Infection Cohort (Balgrist, BASEC 2017-01458). Participants with other infections were assessed in the context of BacVivo (BASEC 2017-02225). All protocols were approved by the local ethics committee.

### Sample collection and processing

During this timeframe, we applied a convenience sampling strategy to acquire and process material from medical procedures of a subset of these patients. We specifically targeted suspected or confirmed staphylococcal infections at the time of the sampling (Suppl. Method S1).

Patient-derived material was homogenized, eukaryotic cells lysed, and antibiotics washed out (Fig. 1A, Suppl. Method S2). Isolated bacteria were then spread-plated on Columbia Sheep Blood agar (CSB, BioMérieux). Absence of growth was interpreted, based on the parallel routine diagnostic, as either true negative, indicating that the infecting strain had likely already been cleared by the treatment or false negative, indicating that it was missed upon sample processing (Fig 1B). Only *S. aureus* mono-cultures were retained for further analysis. Samples CI1140 and CI1150 have been previously described ^12^.

**Figure 1.**
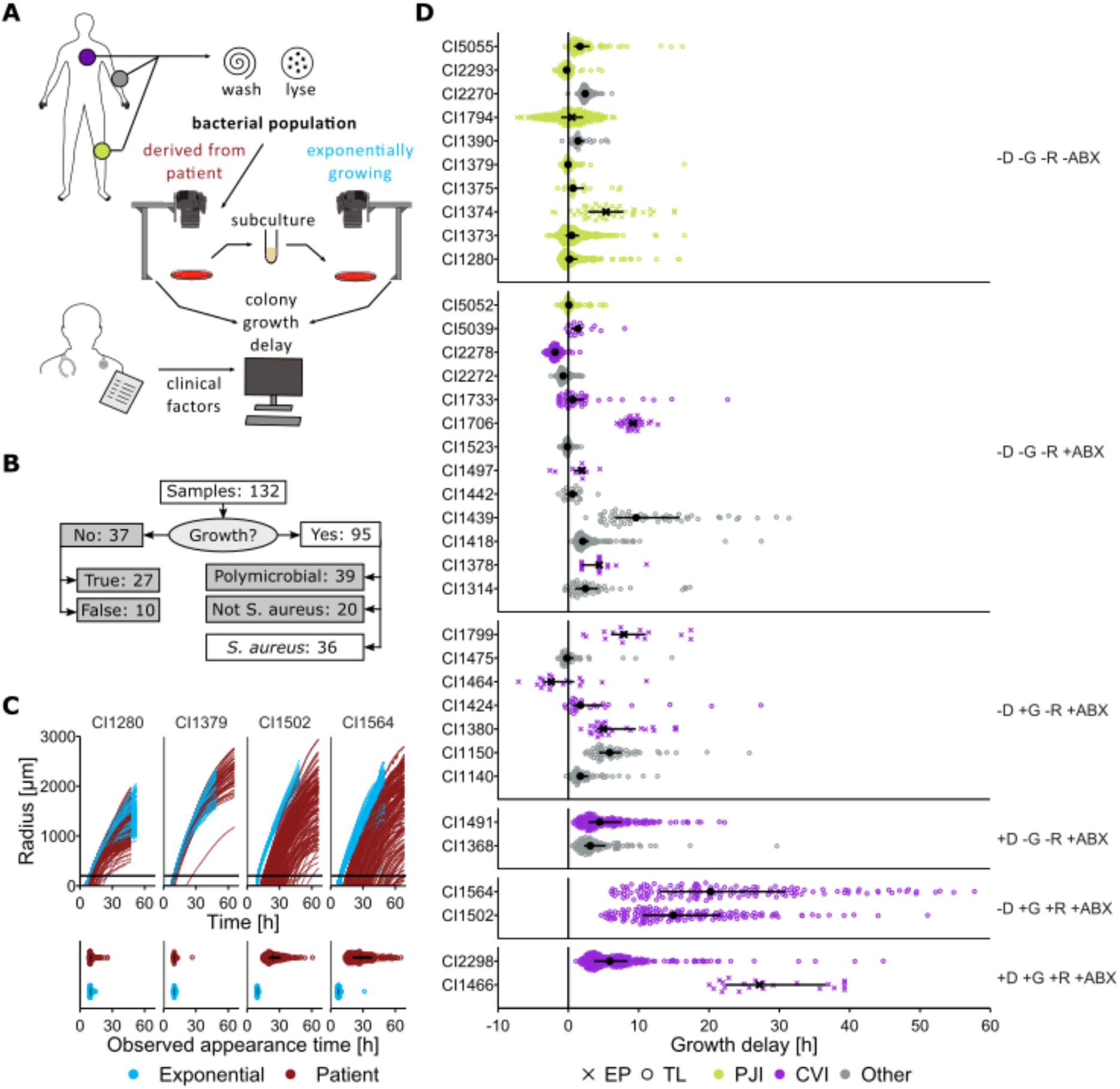
Colony growth-delays of patient-derived *S. aureus*. **A.** Schematic representation of patient sample processing. **B.** Flow-chart representing the study population selection: only *S. aureus* mono-cultures were retained for further analyses. **C.** Top: radial growth curves of colonies formed by patient-derived or exponential-phase bacterial populations (red and blue respectively) from four selected clinical isolates. The black horizontal line corresponds to the threshold of 200 μm for appearance-time determination (Suppl. Method S4). Bottom: corresponding extracted appearance-time distributions. Each dot represents one colony. Interquartile range (IQR) and median are shown in black. **D.** Growth-delay distributions of the 36 patient-derived *S. aureus*. IQR and median are shown in black. Each dot represents one colony, color reflects the clinical category of the infection (PJI, Prosthetic joint infection; CVI, cardiovascular infection) and symbol shape indicates the imaging method used for appearance-time determination (TL, time-lapse images: 26; EP, Endpoint images: 9). Populations are grouped by antibiotic treatment prior to sampling (D, daptomycin; G, gentamicin; R, rifampicin; ABX, any other antibiotic class, including beta-lactams, vancomycin, clarithromycin, metronidazole, ciprofloxacin, levofloxacin, tigecycline or tobramycin. Suppl. Tables S3 and S4)

### Imaging of bacterial colonies

To monitor growth of bacterial colonies, CSB plates were incubated at 37° C and imaged automatically every ten minutes, using a previously described time-lapse (TL) setup.^19^ In a few cases, single-timepoint images (endpoint images, EP), were manually acquired. For most samples, a set of images of one plate on which 20 to 250 colonies had grown was selected for analysis (Suppl. Method S3).

### Colony appearance-time and growth-delay definition

Colony appearance-time was derived from the acquired images using ColTapp (Suppl. Method S4).^19^ Subsequently, growth-delay distributions were obtained by subtracting from these raw appearance-time distributions the baseline appearance-time of the corresponding *S. aureus* clinical isolate, obtained upon subcultivation in tryptic soy broth (TSB, BD) followed by plating while in exponential growth phase (Suppl. Fig. S1 and S2, Suppl. Method S4).

### Biofilm assay

*S. aureus* cultures were grown statically at 37° C in TSB supplemented with 0·15% glucose in 96-well microplates. After 24 h, the supernatant was replaced with fresh TSB supplemented with 0·15% glucose and 10x or 100x minimum inhibitory concentration (MIC) antibiotics or phosphate-buffered saline (PBS) (MIC determination: Suppl. Method S5, Suppl. Table S1). After 24 h, antibiotics were washed out. Viable bacterial load, i.e., number of colony-forming-units per milliliter (CFU/ml), and colony appearance-time of the bacterial population derived from the biofilm or the entire static-stationary culture (including biofilm and its supernatant) was quantified by spread-plating on CSB agar (Suppl. Method S6, Suppl. Fig. S3). Additionally, proportion of rifampicin resistant mutants was assessed using TSB agar supplemented with 100x MIC RIF.

### Antibiotic persister assay

To determine persister levels, killing and regrowth kinetics in liquid medium were examined. Bacterial populations derived from the biofilm assay were diluted to an aimed inoculum of 2 × 10^5^ CFU/ml in TSB, supplemented with either 40x MIC flucloxacillin or PBS. These parallel cultures were incubated at 37° C, shaking at 220 rpm. Viable bacterial load was monitored over time by subsampling the cultures, washing out antibiotics, and spot-plating serial dilutions.

### Statistical analysis

All statistical analyses and plots were performed using R 4·0·3, R Studio and ggplot2.^20^ The effect of clinical parameters on the median growth-delay of bacteria isolated from patients was assessed with univariable and multivariable linear regressions. Dunnett tests were used to compare *in vitro* antibiotic treatments with the corresponding control. Specific pairwise comparisons of *in vitro* assays were computed from linear regression analysis with interaction terms followed by estimated marginal means *post-hoc* tests (multivariate t-distribution based p-value correction, emmeans package).^21^

### Data availability

Detailed methods and additional figures/tables are provided in the Supplementary Material. Raw data, including images and data resulting from image analyses will be available upon publication on Figshare.

## Results

### Clinical isolates collection

We collected a total of 132 samples from 107 patients with difficult-to-treat infections. To quantify colony growth-delay as a proxy for dormancy depth, we isolated bacteria from these patient samples and plated them on nutrient-rich agar (Fig 1A). Out of the 95 samples yielding growth, 36 samples from 30 patients grew *S. aureus* mono-cultures (Fig. 1B). Cultures of other or multiple species were excluded. These 36 samples consisted of native and prosthetic heart valves, vascular grafts, pacemakers, prosthetic joints, bronchoalveolar lavages, necrotic tissue or pus. They had been recovered from cardiovascular infections (CVI, n = 15, 41·7%), prosthetic joint infections (PJI, n = 9, 25·0%) or other clinical categories of infections (n = 12, 33·3%) (Suppl. Table S2).

### Patient-derived *S. aureus* exhibit heterogenous colony growth-delays

Colonies resulting from *S. aureus* plated directly after sampling exhibited heterogeneous appearance-times, which led to colony size heterogeneity (Fig. 1C and 1D). The degree of appearance-time heterogeneity varied substantially between bacterial populations isolated from different patient samples.

Subculturing of each clinical isolate in nutrient-rich medium and plating it from exponential growth phase resulted in reduction of appearance-time heterogeneity (with IQR of appearance time distributions of 0·5-18 h for patient-derived populations and 0·2-2 h for exponential-phase bacterial populations Fig. 1C, Suppl. Fig. S1 and S2). The median time required for exponential-phase bacteria to appear on the plate was considered as time needed for actively dividing colony-forming-units to become observable colonies based on the corresponding strain properties.^19^ Delays from this baseline appearance-time reflected the environmentally-induced phenotypic state of the bacteria. In some cases, most colonies exhibited marginal growth-delays (e.g., CI1280, CI1379, Fig. 1C), suggesting that most bacteria recovered from the infection site were actively dividing within patients. In other cases, extended growth-delays of up to 57·6 h were observed (e.g., CI1502, CI1564, Fig. 1C), indicating dormant states within patients.

Extreme growth-delays co-occurred with the highest variance (Fig. 1D) and were accompanied by a global increase of delay for the entire population. Previously, we summarized colony growth-delay distributions by quantifying their tail, with an absolute threshold based on radius or appearance-time.^12,15,22^ Here, given the correlation of the median of the distributions with the proportion of colonies appearing later than six hours (Suppl. Fig. S4), we used the median growth-delay as an estimator of population-wide dormancy.

Since antibiotic therapy and clinical category of infection appeared as promising predictors of phenotypic heterogeneity (Fig. 1D), we subsequently investigated the effect of these and other clinical parameters on median growth-delay.

### Growth-delay of patient-derived *S. aureus* is associated with antibiotic treatment regimens

To explore which clinical parameters predicted patient-derived *S. aureus* median growth-delay, we used a combination of univariable and multivariable linear regression (Fig. 2, Suppl. Table S3 and S4). We included general patient and infection characteristics as well as characteristics of the within-patient environment encountered by bacteria as predictors. The latter included antibiotic treatment of the patient any time prior to sampling, omitting the complexity of time dynamics. The five most prescribed antibiotics in this study were included as individual factors.

**Figure 2.**
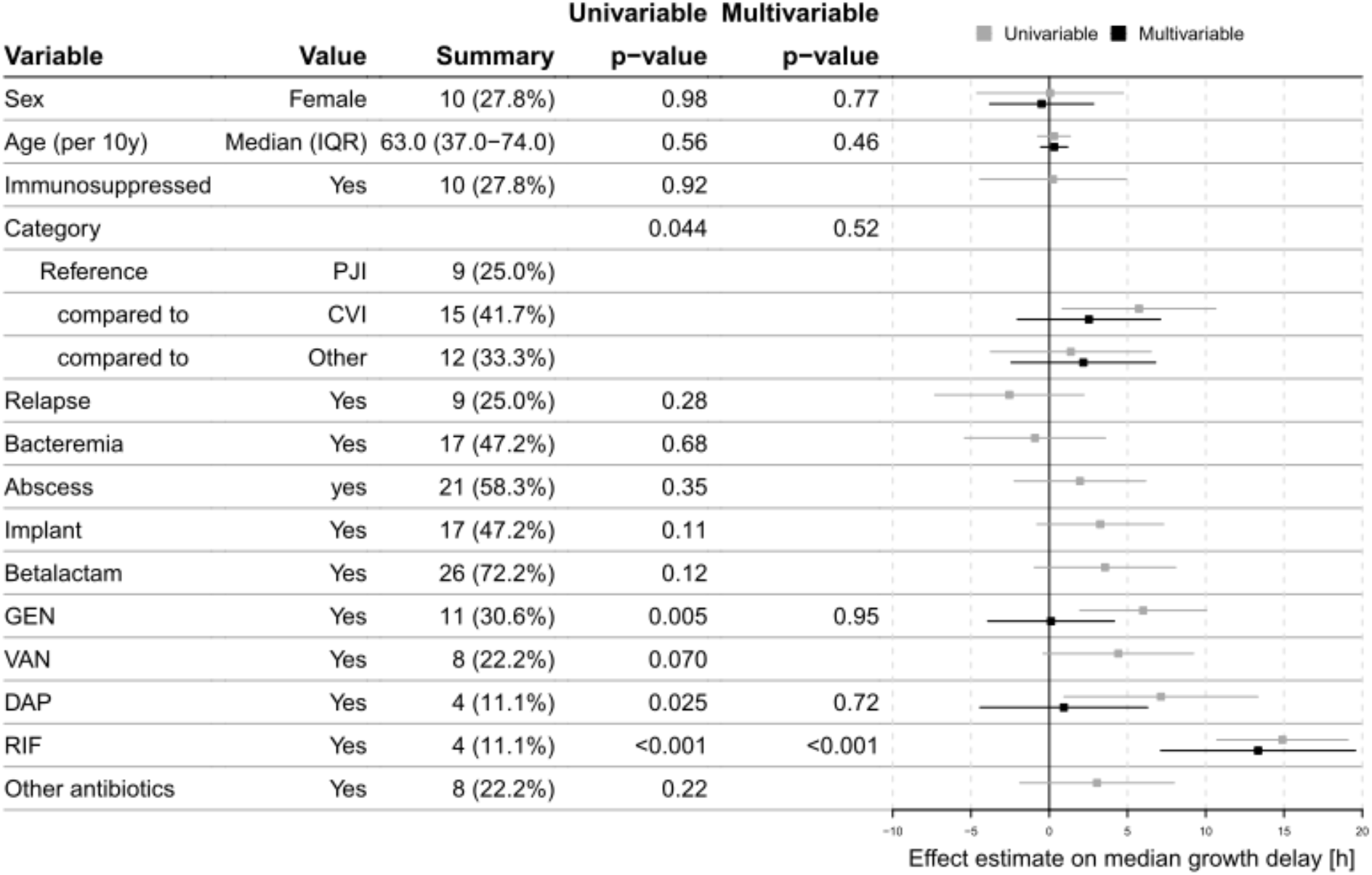
Effect of 14 clinical parameters on the median growth-delay of patient-derived *S. aureus* (n = 36), based on univariable and multivariable linear regression. Sex, age, and parameters with a p-value below 0.05 in the univariable model were included in the multivariable analysis. Categorical and continuous parameters are summarized with count (percentage) for the level indicated or median (interquartile range, IQR), respectively. For the factor clinical category, PJI (prosthetic joint infections) was used as the reference level to which CVI (cardiovascular infections) and other infections were compared to. GEN: gentamicin, VAN: vancomycin, DAP: daptomycin, RIF: rifampicin. “Other antibiotics” includes clarithromycin, metronidazole, ciprofloxacin, levofloxacin, tigecycline or tobramycin.

After multivariable adjustment, rifampicin treatment (RIF) was significantly associated with larger median growth-delay (mean and confidence interval (CI): 13·3 h [7·13, 19·6], respectively).

Although certain explanatory variables were correlated due to inherent differences in types of infection and associated standard-of-care (Suppl. Fig. S5), the direction, effect size and significance of RIF was robust when additionally adjusting for technical variables, i.e., imaging method and preparation delays, or when subsampling the data, i.e., excluding EP imaged samples, multiple samples from the same patient except the latest, or PJI samples (Suppl. Fig. S6).

When considering the percentage of colonies with growth-delay larger than six hours rather than the median as outcome, we obtained comparable effects for RIF (mean and CI: 52·0% [20·3, 83·8]). Additionally, in this assessment, vancomycin treatment (VAN) was significantly associated with an increased percentage of colonies with large growth-delays (31·1% [7·84, 54·4], Suppl. Fig. S7).

### Biofilm-embedded *S. aureus* surviving high concentrations of rifampicin exhibit increased colony growth-delays

Based on the clinical observations, we sought to evaluate the effect of antibiotic exposure on colony growth-delay of *S. aureus* derived from a heterogeneous environment. To mimic this environment *in vitro*, we grew static biofilms with a subset of clinical isolates and exposed them to five routinely used antibiotics with different modes of action (FLX, clindamycin (CLI), gentamicin (GEN), levofloxacin (LVX) and RIF) at 10x and 100x MIC (Suppl. Method S5, Suppl. Table S1).

Bacterial populations derived from biofilms which had been exposed to FLX, CLI, GEN and LVX exhibited colony growth-delay distributions similar to those of the corresponding no-antibiotic control. In contrast, populations derived from biofilms exposed to RIF displayed increased growth-delay (Fig. 3A, Suppl. Fig. S8).

**Figure 3.**
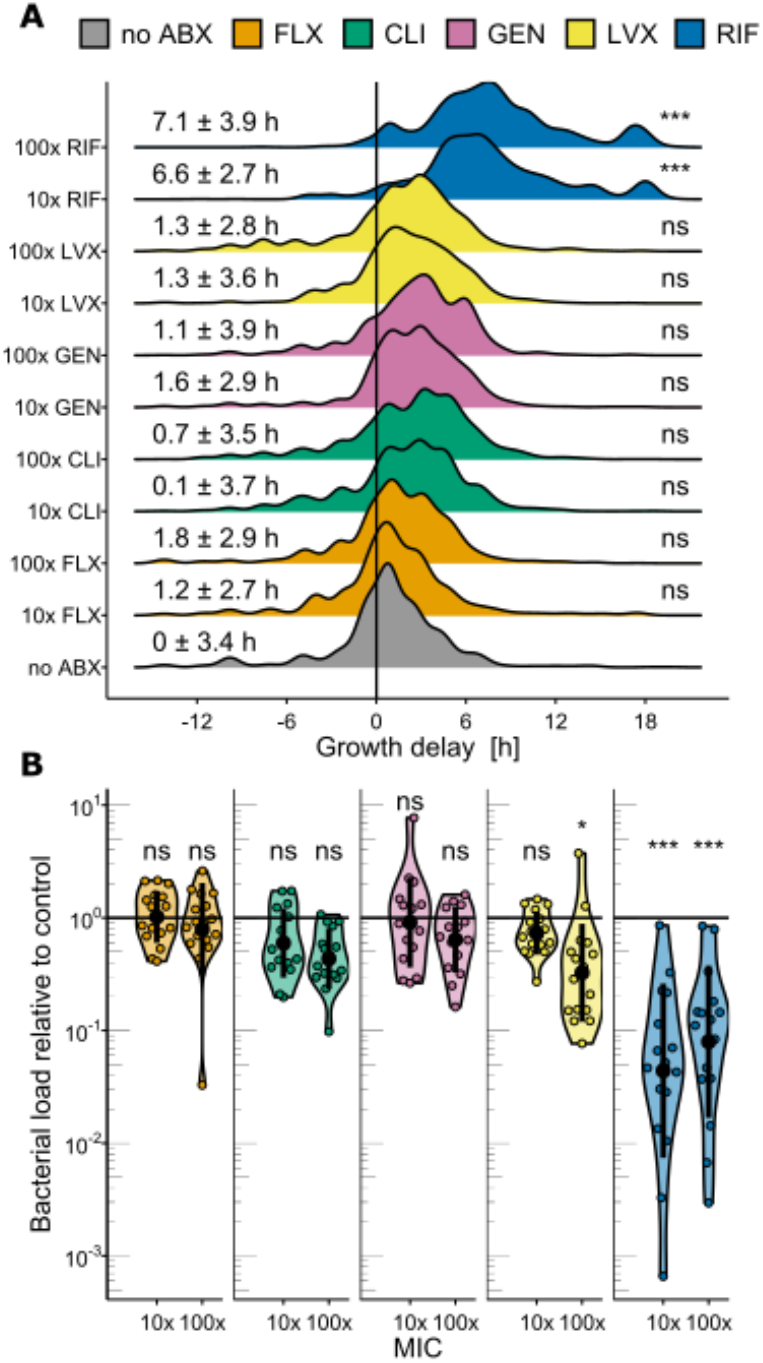
Biofilm assay: screening of 17 representative clinical isolates. **A.** Growth-delay distributions from biofilm-embedded *S. aureus* populations exposed to either a no antibiotic control (no ABX) or flucloxacillin (FLX), clindamycin (CLI), gentamicin (GEN), levofloxacin (LVX) or rifampicin (RIF) at 10x and 100x minimum inhibitory concentration (MIC), combining all clinical isolates. Mean and standard deviation of median growth-delay per clinical isolate are given on the left and significance of corresponding statistical tests is indicated on the right. **B.** Biofilm eradication efficacy measured as bacterial load recovered relative to the no-antibiotic control bacterial load. Each dot represents one clinical isolate. Black dots and bars represent mean and standard deviation. ns: non-significant; * p < 0.05; *** p < 0.0001 based on a Dunnett’s test comparing all antibiotic and concentration combinations to the no-antibiotic control.

We found that the bacterial load of the biofilm (4·9 ± 1·9 × 10^8^ CFU/ml, for 17 clinical isolates’ no-antibiotic control), was not reduced by exposure to most antibiotics, except for the highest LVX concentration (3·1 ± 5·4 × 10^8^ CFU/ml for 100x MIC), and both concentrations of RIF (5·3 ± 6·4 × 10^7^ CFU/ml and 9·1 ± 13 × 10^7^ CFU/ml for 10x and 100x MIC, respectively, Fig. 3B).

Overall, our data showed that RIF efficiently killed biofilm-embedded *S. aureus*, but surviving bacteria exhibited an increased delay in growth resumption. However, we did not observe a correlation between median growth-delay and bacterial load according to treatment (Suppl. Fig. S9).

### Increased growth-delays result in antibiotic tolerance

Next, we investigated whether delays in growth resumption resulting from RIF treatment promote antibiotic tolerance. Therefore, we challenged bacterial population derived from the biofilm with 40x MIC of the beta-lactam FLX in liquid nutrient-rich medium and monitored survival over time. For this experiment, we included four clinical isolates that displayed various levels of growth-delay *ex vivo* (Fig. 1C). Moreover, since RIF is not used as a monotherapy due to the high rate of resistance emergence,^23^ we additionally included combination treatments of RIF with CLI or LVX.

Any treatment containing RIF reduced the bacterial load significantly (Fig. 4A). Combination of RIF with CLI marginally reduced the killing capacity of RIF alone while combination with LVX resulted in slightly improved killing. As expected, the proportion of resistant mutants was high upon RIF monotreatment (0·053 ± 0·15%) but reduced – often below our detection threshold – if RIF was combined with CLI or LVX (Fig. 4A, Suppl. Method S7). Regarding colony growth-delay distributions, RIF combination treatments had comparable results as RIF mono treatment – i.e., a global shift and longer tail as compared to the no antibiotic control – with slight strain-dependent deviations in magnitude (Fig. 4B, Suppl. Fig. S10).

**Figure 4.**
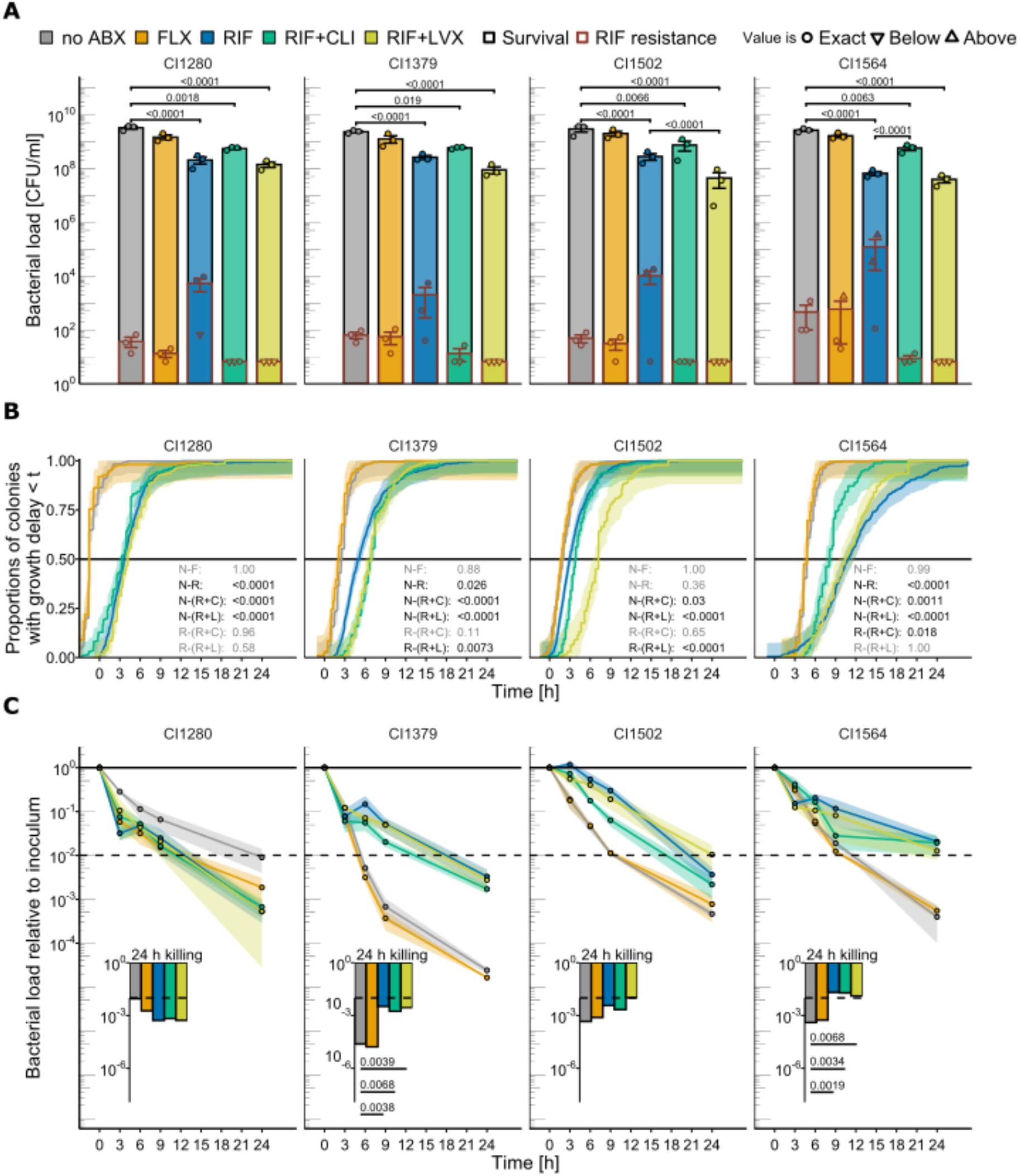
Biofilm assay followed by persister assay for four clinical isolates **A.** Bacterial load (CFU/ml) of the biofilms for the no-antibiotic control (no ABX) or after exposure to flucloxacillin (FLX), rifampicin (RIF), the combination of rifampicin and clindamycin (RIF + CLI) or rifampicin and levofloxacin (RIF + LVX) (black contour) and corresponding load (CFU/ml) of RIF resistant mutants (overlaid, with a red contour). Mean and standard deviation are shown. Dots represent biological replicates (n = 3), and shapes indicate if the value is exact or below/above our detection range (Suppl. Method S7). **B.** Empirical cumulative distribution function of colony growth-delay based on the three biological replicates combined. Shaded area depicts the confidence interval. The black line allows visual extrapolation of median growth-delay for each condition. N, no ABX; F, FLX; R, RIF; R+C, RIF+CLI; R+L, RIF+LVX. All pairwise comparisons performed are indicated and displayed with corresponding p-value in black or grey if significant or non-significant, respectively (e.g., N-F stands for median growth-delay of the no-antibiotic control versus that of the FLX exposed biofilm and is always shown in grey because non-significant). **C.** Time-kill curve upon 40x MIC FLX challenge in liquid medium. Dots and shaded area represent mean and standard error of three biological replicates. The dashed line labels a 99% reduction of the initial bacterial load, to allow visual extrapolation of the minimal duration to kill 99% (MDK99). Bacterial load (A), median growth-delay (B) and survival after 24 h 40x MIC FLX challenge (C) were assessed with linear regressions with interaction terms followed by pairwise comparisons computed with estimated marginal means *post-hoc* tests (p-value correction based on multivariate *t-*distribution).

When inoculating an equivalent bacterial load (2 × 10^5^ CFU/ml) from these pre-exposed bacteria in liquid nutrient-rich medium without antibiotics, similar regrowth dynamics were observed: any treatment containing RIF resulted in prolonged recovery periods of up to 9 h (Suppl. Fig. S11). In some cases, the bacterial load further dropped during this recovery period, potentially reflecting death due to post-antibiotic stress.^24^

These regrowth kinetics were mirrored by the killing kinetics in the parallel liquid culture that had been supplemented with 40x MIC FLX: treatments including RIF resulted in longer time to kill the same fraction of the population (Fig. 4C). This resulted in higher rates of bacterial survival after 24 h, for all but one of the clinical isolates (CI1280). Generally, reproducible but distinct kinetics were observed, and bacterial load reduction differed between clinical isolates. For example, CI1379 and CI1564 both displayed classical biphasic killing curves, but the latter had an inherent higher ability to persist in FLX than the former.

In conclusion, exposing *S. aureus* biofilms to RIF mono and combination treatments effectively reduced viable bacterial load. However, any treatment containing RIF resulted in increased growth-delays, which in three of the four tested clinical isolates correlated with increased antibiotic tolerance (Suppl. Fig. S12).

## Discussion

Colony size heterogeneity has been repeatedly linked to difficult-to-treat infections and attributed to the presence of persisters *in vivo*.^9–12^ Yet, there is currently a lack of systematic comparative analysis quantifying this phenomenon across clinical presentations.

In this study, we quantified within-patient *S. aureus* phenotypic heterogeneity in difficult-to-treat acute bacterial infections, by monitoring colony growth dynamics directly upon sampling from patients. Different bacterial populations exhibited substantially different degrees of growth-delay heterogeneity. Those exhibiting the widest heterogeneity and the highest median growth-delays were recovered from cardiovascular infections (CVI) treated with multiple antibiotics. Upon multivariable adjusted linear regression, RIF treatment prior to sampling was significantly associated with increased median growth-delay.

Following our observation that antibiotic treatment generates a detectable signal on within-patient bacterial phenotypic heterogeneity, future studies with a more controlled sampling design will be necessary to elucidate specific effects of patient treatment and underlying disease. Indeed, unavoidable biases were introduced by the convenience sampling design of this study. Most importantly, standard of care for prosthetic joint infections (PJI) and CVI differed substantially: surgical procedures are part of gold-standard care for PJI and more rarely performed to treat CVI. In our collection, CVI sampled were mostly severe life-threatening cases, which had been treated with combination regimens at the time of surgery. In contrast, most patients with PJI had not been treated with antibiotics prior to surgery. Another limitation of our analysis was to consider antibiotic prescriptions prior to sampling regardless of treatment duration and last occurrence of antimicrobial drug intake. We speculate that the phenotype observed is likely to be affected by temporal dynamics and to be subject to drug interactions instead of the result of additive effects.

Nevertheless, with our *in vitro* experiment, we demonstrated the biological validity of the link between RIF treatment and increased growth-delays, identified upon statistical analysis of our clinical data. Concurrently, the performance of RIF in reducing bacterial load was superior to other tested antibiotics, consistent with previous studies on staphylococcal biofilm eradication.^25,26^ Co-occurrence of decreased bacterial load with increased median growth-delays hinders from elucidating whether RIF treatment does induce longer growth-delays, or simply selects a pre-existing subpopulation with long growth-delays by killing the bulk of the population with short growth-delays. Previous literature indicates that antibiotics, including RIF, act as an environmental stressor that can induce persister formation.^16,27,28^ In the context of this study, selective killing of the bulk of the population with shortest growth-delays could suffice to explain our observation of unimodal growth-delay distributions with higher median and variance after biofilm treatment with RIF. The observation of such unimodal rather than bimodal distributions is consistent with previous studies looking at isogenic populations upon stress exposure.^15,29,30^ More research is needed to disentangle population kinetics linked with persister selection and induction. Furthermore, the growth-delay of some populations derived from RIF-treated-patients was more extreme than delays observed upon any antibiotic exposure *in vitro*. This discrepancy could be attributed to additional host stressors and/or different population kinetics within-patient.

In conclusion, we demonstrated that antibiotic treatment was the major factor associated with growth-delay, which was linked with antibiotic tolerance *in vitro*. Future studies dissecting clinical factors and population kinetics will be necessary to identify the larger clinical relevance of this finding. Given that growth-delay heterogeneity is an indicator for presence of antibiotic tolerant bacteria and different levels of heterogeneity can be observed *ex vivo*, introduction of colony growth dynamics monitoring in microbiological diagnostic routine is important. In parallel, development of new therapies specifically targeting persisters will be crucial.

## Author contributions

SDB, RAS, RDK, BH, ASZ designed the study; BH, ASZ coordinated ENVALVE, VASGRA and BACVIVO cohorts; YA coordinated the PJI cohort; YA, PZ, CAM, SDB, RAS, ASZ acquired patient samples; NE, YA, SDB, BH collected clinical data; JB, SMS, MH, TAS, AGM processed the patient samples; JB, MB, SMS, CV, MH, TAS performed experiments; JB, MB, SMS, CV designed and interpreted experiments; JB, MB, CV performed image analysis; JB, MB, RDK performed statistical analysis; JB, MB wrote the first draft of the manuscript; SMS, CV, SDB, RDK, BH, AZ critically revised the manuscript.

## Acknowledgements

We are grateful to our patients for their participation in the study and the study nurses Caroline Mueller and Simone Bürgin for their excellent work. We also thank Christine Laich and Christine Vögtli for administrative assistance, and the technicians of the Institute of Medical Microbiology of the University of Zurich for their expert help and assistance. We thank the students Vera Beusch, Milos Duknic for assistance in preliminary data analysis.

