## Supplementary material for "Quantification of within-patient *Staphylococcus aureus* phenotypic heterogeneity as a proxy for presence of persisters across clinical presentations"

### Supplementary Methods

#### Suppl. Method S1: Patient samples collection

Bacteria were sampled from patients with a suspected or confirmed staphylococcal infection. Often, the etiological agent was not confirmed at the time of patient material processing and colony growth imaging, since species identification was performed in parallel by the routine clinical microbiology laboratory.

If colony morphology on imaged agar plates indicated polymicrobial cultures, contaminations, or a different species than that identified by the diagnostic laboratory, additional colonies were screened using a matrix-assisted laser desorption/ionization time-of-flight (MALDI-ToF) mass spectrometer and the MALDI BioTyper software (BRUKKER).

Patient samples consisted of various materials: explanted prosthesis or sonication fluid from prosthetic joint infections (PJI); debrided tissue, explanted cardiac devices and one blood culture from cardiovascular infections (CVI); pus, skin or tissue samples from soft tissue, skin, and abscess related infections; nasal swabs or bronchoalveolar lavages from respiratory tract infections.

#### Suppl. Method S2: Patient samples processing protocol

Tissue samples and prosthetic heart valves were cut in small pieces, weighted and phosphate-buffered saline (PBS) was added (two times the weight of the sample). The sample was then homogenized with metal beads in a 2 ml Eppendorf tube with a TissueLyser (Qiagen) for 10 min at a frequency of 30'000 Hz. The tube was centrifuged for 5 min at 1'500 rpm and the supernatant collected. The remaining tissue pellet was resuspended in 0.5 to 1 ml water to lyse the eukaryotic cells and centrifuged again with the same settings, supernatant collected and pooled with the previous collection. The pooled supernatant was then washed twice with water by centrifugation at 14'000 rpm for 3 min. The pellet was resuspended in water and serially diluted. Multiple dilutions were spread-plated on Columbia Sheep Blood agar plates (CSB, BioMérieux).

Prosthetic joints, pacemakers, cables, vascular grafts, and other foreign body material was sonicated for 5 to 10 min in 50 ml PBS. Sonicated liquid was collected, centrifugated for 10 min at 4'000 rpm and the supernatant discarded. Depending on the sample type, a second sonication step of the material in 50ml fresh PBS was done. The pellets were resuspended in 10 ml of PBS each, pooled together and centrifugated with the same settings as before. Supernatant was again discarded, the pellet resuspended in 1 ml water and transferred to a 2 ml Eppendorf tube. The sample was washed twice with water by a 3 min centrifugation at 14'000 rpm. Then, the remaining pellet was resuspended in 0.5 ml water and multiple dilutions were spread-plated on CSB agar plates.

#### Suppl. Method S3: Colony density on imaged agar plates

To avoid a bias resulting from differences in colony density on the plates (i.e., number of colonies per plate), a set of images of one plate on which 20 to 250 colonies had grown was selected for analysis, for most clinical isolates. There were eight exceptions to this rule: for three clinical isolates, images with more than 250 colonies were analyzed (CI1280, n = 253; CI1373, n = 368; CI1794, n = 272), either because images of the optimal dilution were affected by technical problems or because the optimal dilution was not captured due to an unexpectedly high bacterial load. For five clinical isolates (CI1378, CI1464, CI1497, CI1799, CI5039), images with less than 20 colonies were analyzed. Four of these cases consisted in endpoint images and of these, three (CI1378, CI1464,

CI1799) had several plates imaged. In these three cases, the data extracted from up to four plates with each up to 16 colonies were pooled.

**Suppl. Method S4: Colony appearance-time definition and calibration**

Colony appearance-time, defined as the time to reach a radius of 200 µm, was either directly derived from the colony radial growth curves generated by TL imaging or estimated by linear extrapolation based on colony radii measured on EP images. EP images were usually acquired after 24 h of incubation and the appearance-time of colonies that appeared after this timepoint was set to the imaged timepoint (i.e., 24 h).

For calibration, each *S. aureus* clinical isolate was subcultured in tryptic soy broth (TSB) overnight (o/n) at 37° C, shaking at 220 rpm. The culture was brought to exponential phase by allowing a 1:10 dilution to regrow for 2 h, and then serial dilutions were plated on CSB agar for TL imaging. The median appearance-time of this culture was defined as the baseline appearance-time and subtracted from the appearance-time of each colony deriving from the same clinical isolate (*ex vivo* as well as *in vitro*) to define a growth-delay measurement attributable to the growth conditions.

**Suppl. Method S5: Antibiotic susceptibility evaluation**

For each clinical isolate, the minimum inhibitory concentration (MIC) of five antibiotics (oxacillin (OXA), gentamicin (GEN), clindamycin (CLI), levofloxacin (LVX), rifampicin (RIF)) was assessed with ETEST strips (BioMérieux). The strips were placed on Mueller-Hinton agar plates that had been fully inoculated with a cotton swab dipped into a 0.1 McFarland suspension. The inhibition zone was visually inspected after 24 h incubation at 37° C. For *in vitro* experiments, reference MIC values were determined (Suppl. Table S1).

**Suppl. Method S6: Biofilm assay**

*S. aureus* o/n cultures were diluted to an optical density ($OD_{600nm}$) of 0.0025 in TSB supplemented with 0.15% glucose, 200 µl added per well to 96-well Nunclon Delta-Treated microplates (Thermofisher) and grown statically at 37° C for 24 h. The formed biofilms were subsequently exposed for 24 h to the antibiotic treatment, by replacing the supernatant with fresh TSB supplemented with 0.15% glucose and 10x or 100x minimum inhibitory concentration (MIC) antibiotics or phosphate-buffered saline (PBS) (Suppl. Fig. S3A, for MIC determination see Suppl. Method 5, Suppl. Table S1).

For the biofilm assay with single antibiotics exposure (Fig. 3) the wells' content was divided in three parts (Suppl. Fig. S3B1): the supernatant (200 µl), transferred to another microplate; the sedimented but not attached bacterial cells, collected with two gentle rinses with 100 µl PBS and transferred to another microplate; and the bacterial cells embedded in the biofilm matrix, collected by mechanical disruption by vigorous resuspension in 200 µl PBS. Bacterial load within the collected fractions as well as colony growth-delay upon plating were assessed (Suppl. Fig. S3C).

Subsequently, since bacterial load and rifampicin resistance evolution rate were consistent across these three fractions (Suppl. Fig. S13), the entire static stationary culture (including biofilm and supernatant) was processed for the biofilm assay with rifampicin combinations exposure (Fig. 4). Separation steps were omitted and the 200 µl of supernatant were utilized to disrupt the formed biofilm (Suppl. Fig. S3 B2). Bacterial load and proportion of RIF resistant mutants within the entire static-stationary culture as well as colony-growth delay were assessed and subsamples of each population were inoculated in TSB for the persister assay (Suppl. Fig. S3 C, D).

In either protocol, before assessment of the phenotypes following antibiotic exposure, antibiotics were washed out twice from the fractions of interest by centrifugating the microplates for 10 min at 4’000 rpm and replacing 150 µl of supernatant with PBS.

**Suppl. Method S7: Bacterial load detection range**

Based on preliminary experiments in the tested conditions, we plated dilutions of the population of interest aiming for between 20 and 200 colonies per plate, to avoid biases in appearance-time estimation resulting from differences in colony density on the plates. Thus, the bacterial load detection range, which differs between conditions and biological replicates, is the range in which we were able to assess an exact CFU number from the dilutions plated. In most cases, an exact number could be determined but, in some cases, especially for the RIF resistant mutants, less predictable due to stochasticity of mutation events during antibiotic treatment, the plates were either overgrown (above detection range) or less than 10 CFU could be counted (below detection range).

### Supplementary Figures

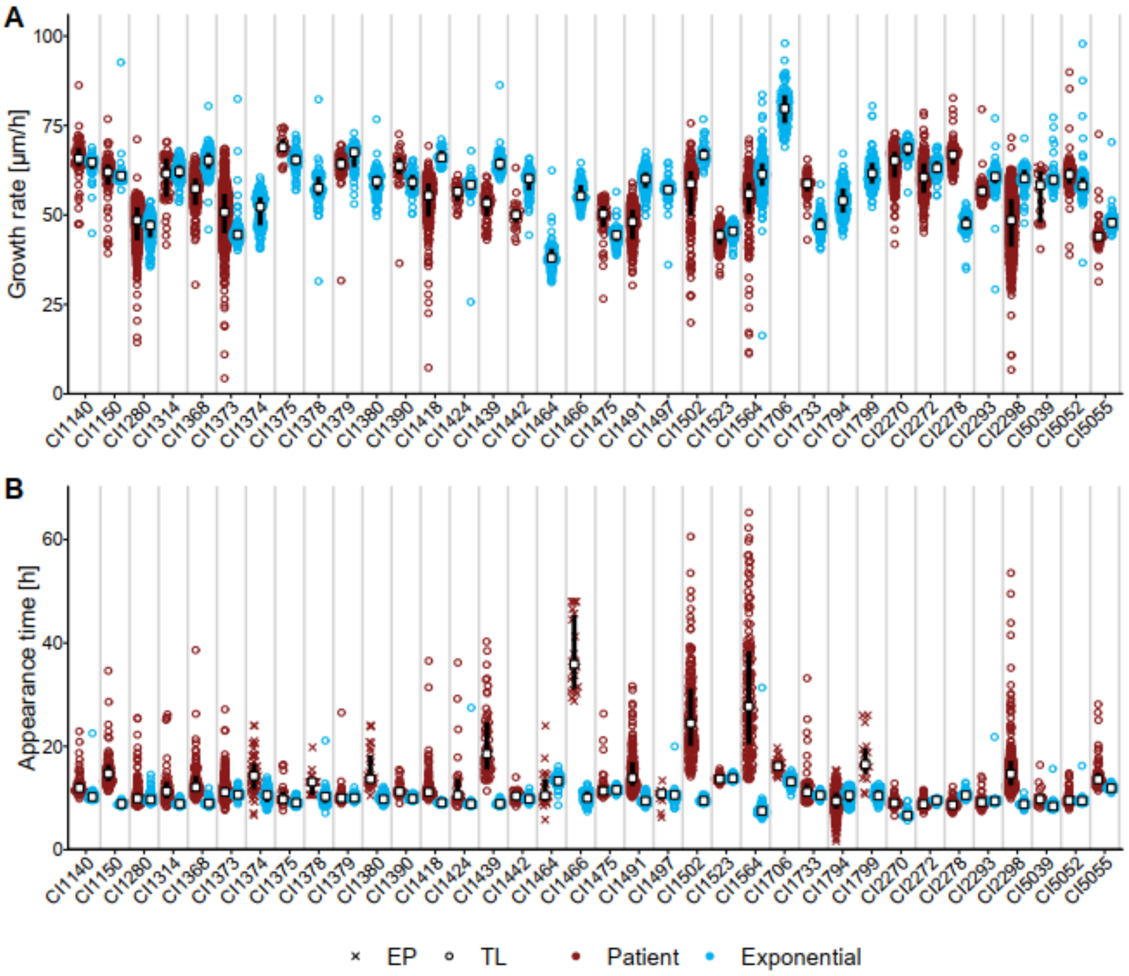

**Suppl. Fig. S1: Overview of colony growth rate and appearance-time** (**A.** and **B.** respectively) of patient-derived (red) or exponential-phase (blue) bacterial populations. Each dot represents one colony, white squares indicate the median values and black bars the IQR. Colony radial growth rate of the EP-imaged patient samples is unknown. The patient samples' appearance-time IQR ranged from 0.5 to 18 h, whereas exponential cultures' appearance-time IQR ranged from 0.2 to 2 h. For each clinical isolate, median values of growth rate and appearance-time from the exponential cultures were used as reference values. Reference growth rate was used to estimate appearance-time of the EP-imaged patient-derived and biofilm-derived bacterial populations, based on colony radius. Reference appearance-time was used to calibrate the appearance-time distributions of patient-derived and biofilm-derived bacterial populations.

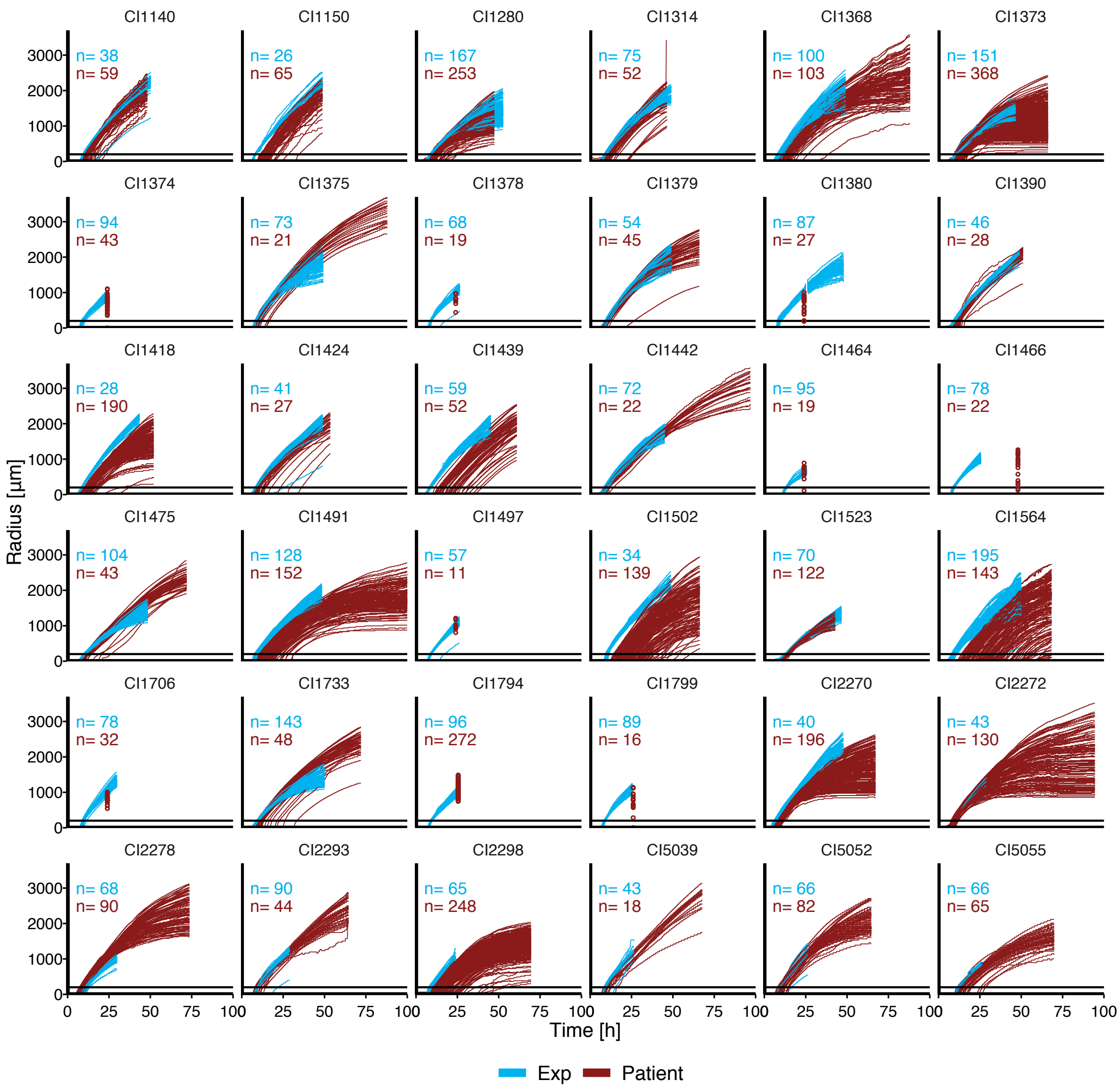

**Suppl. Fig. S2: Colony radial growth curves** of bacterial populations derived from exponential cultures (blue) or directly from patient samples (red). For the patient samples with missing TL data, colony radii measured at an endpoint (EP) are displayed as dots. The horizontal black line indicates the radius threshold of 200 µm for appearance-time determination.

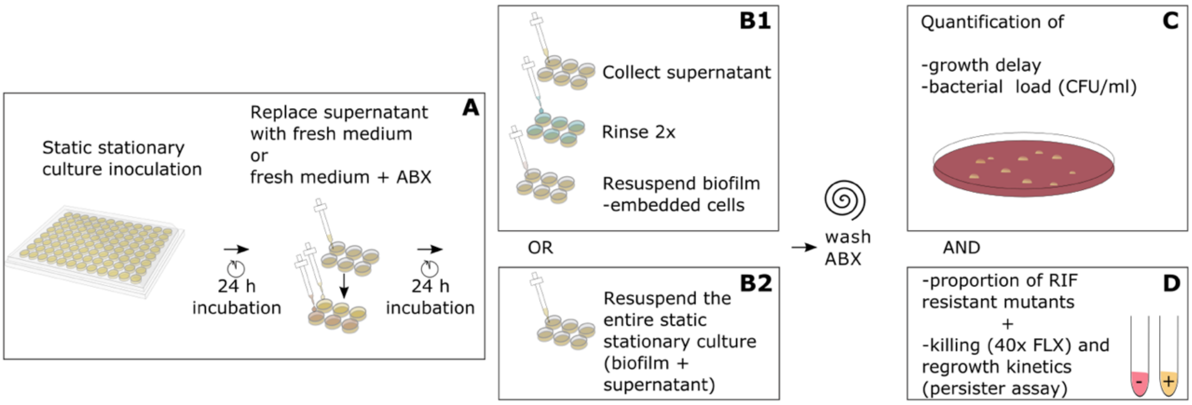

**Suppl. Fig. S3. Schematic representation of the biofilm assay** described in Suppl. Method 6. For Fig. 3: A, B1 and C, for Fig. 4: A, B2, C and D.

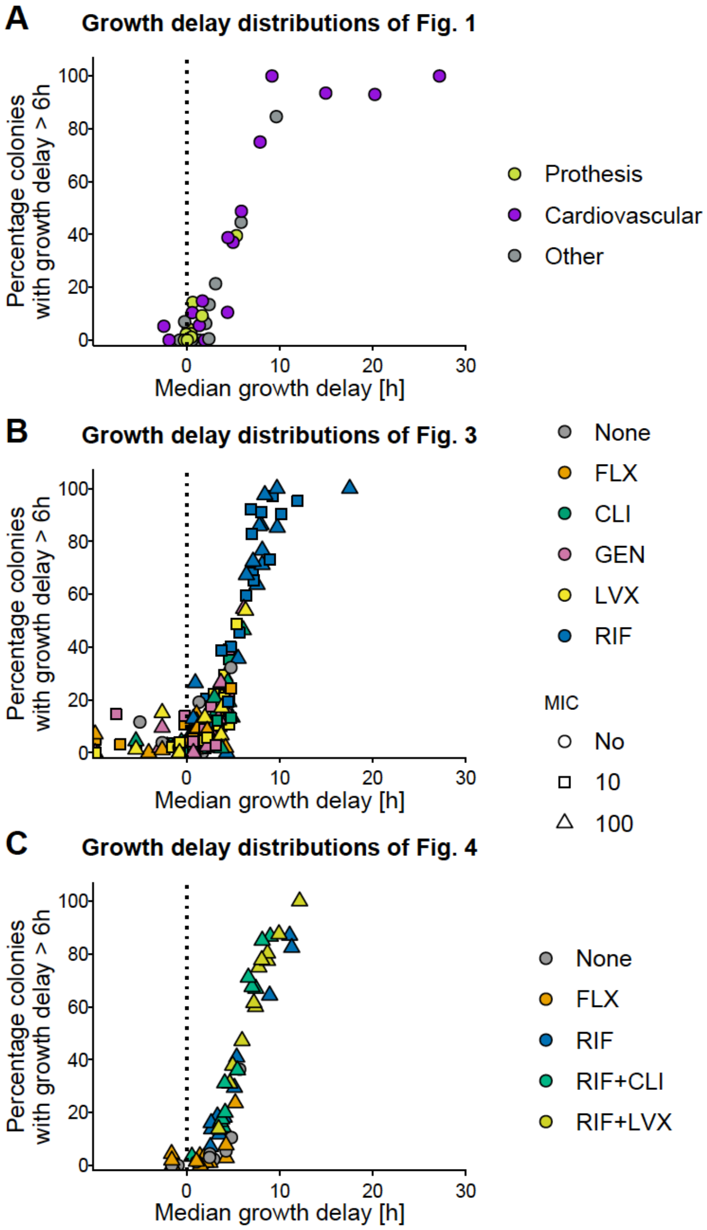

**Suppl. Fig. S4: Relation between summary values of growth-delay distributions.**

For the three sets of growth-delay distributions generated in this study, median of the distribution correlates with percentage of colonies with a growth-delay larger than 6 h.

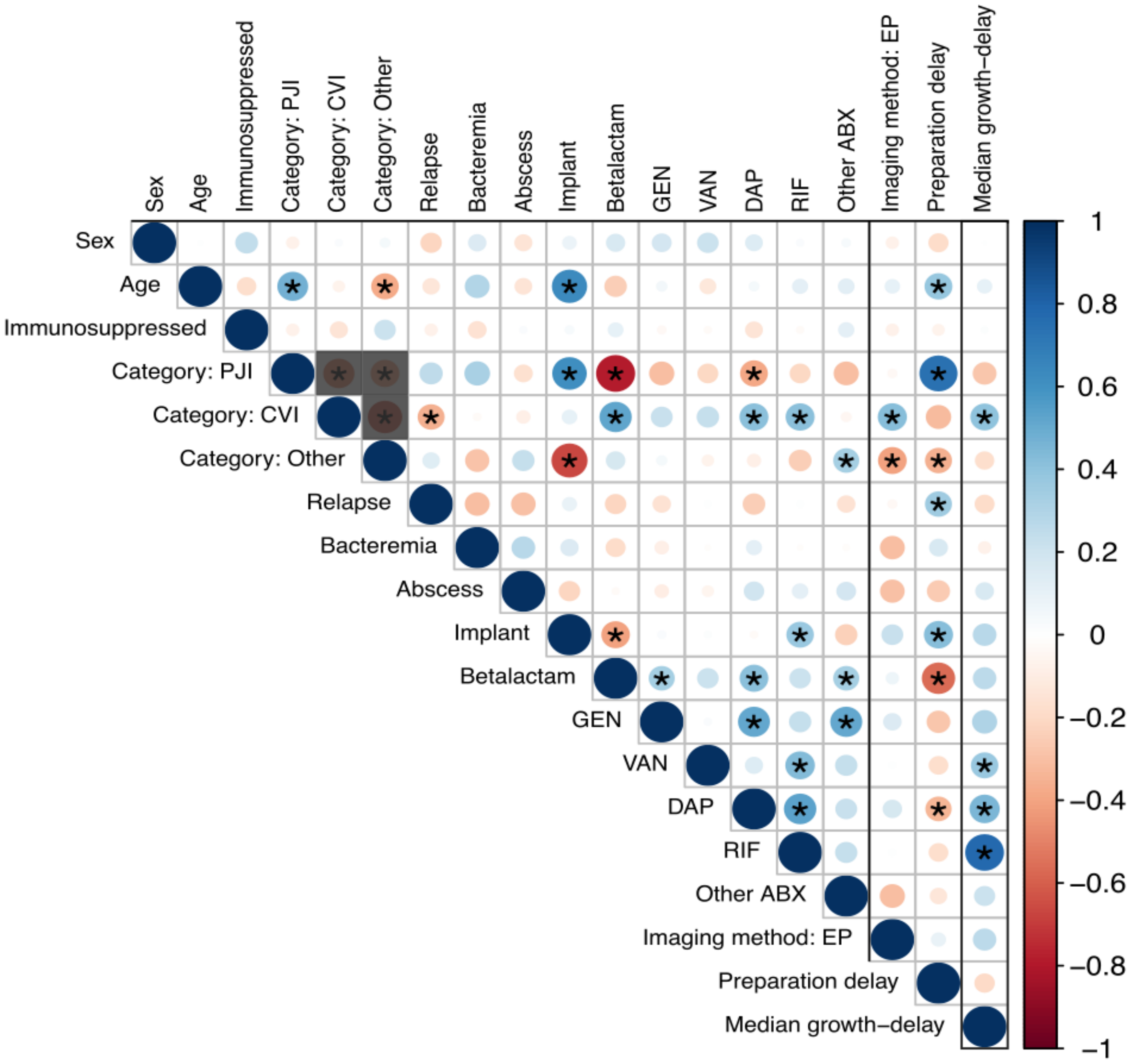

**Suppl. Fig. S5: Pairwise correlation among variables** for all explanatory variables (Suppl. Table S3) and outcome (median growth-delay, last column). The plot was produced with the corrplot R package (https://cran.r-project.org/web/packages/corrplot). The color and size of circles reflect the Pearson correlation coefficient and stars indicate a p-value < 0.05. The grayed squares mask the obvious negative correlation between levels of the variable clinical category of infection (PJI, Prosthetic joint infection; CVI, cardiovascular infection). GEN: gentamicin, VAN: vancomycin, DAP: daptomycin, RIF: rifampicin. Other ABX (other antibiotics) include clarithromycin, metronidazole, ciprofloxacin, levofloxacin, tigecycline or tobramycin.

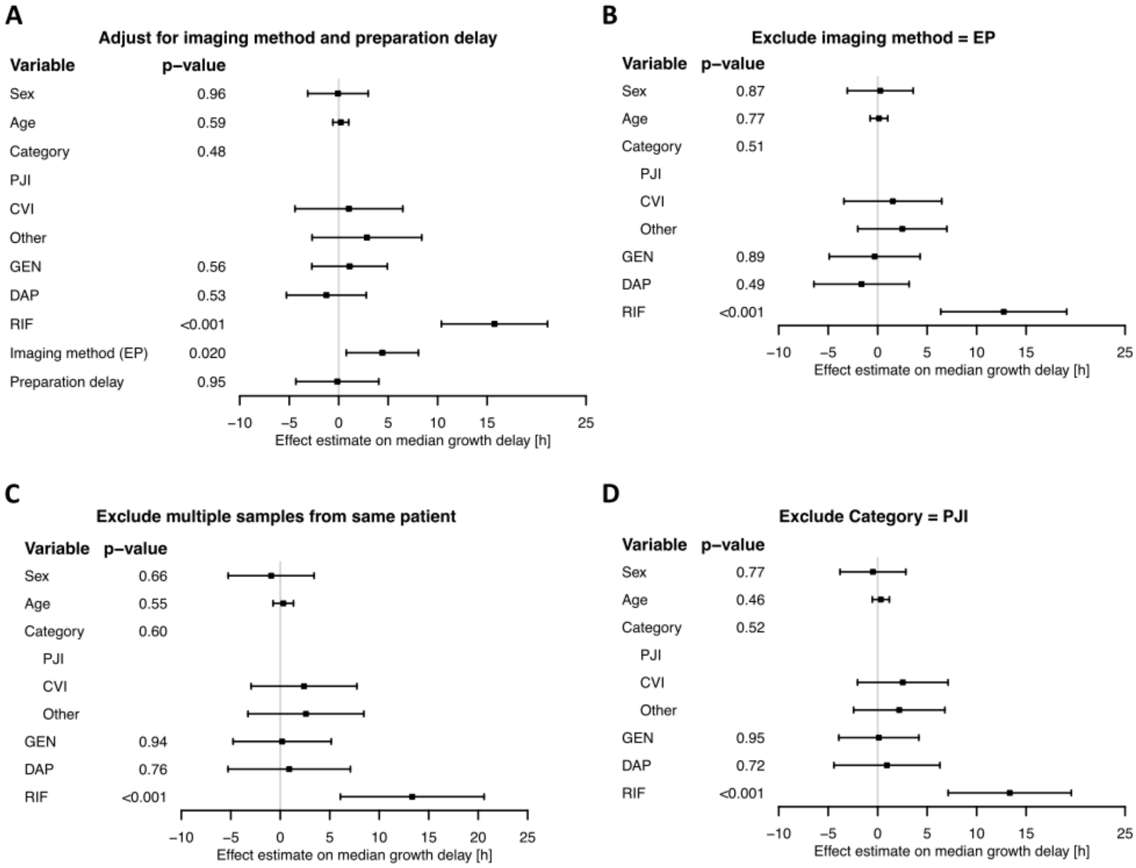

**Suppl. Fig. S6: Sensitivity analysis of the multivariable linear regression** utilized to explain the median growth-delay (Fig. 2). The direction, effect size and significance of the association of rifampicin treatment (RIF) with increased median growth-delay was robust upon:

- **A.** adjustment for two additional technical variables: imaging method and sample preparation delay (Suppl. Table S3, S4),
- **B.** subsampling of the data by excluding all EP imaged samples (which were significantly associated with higher median growth-delay, see **A.**),
- **C.** subsampling the data by excluding multiple samples from the same patient (i.e., including only the latest sample for the five patients from whom more than one sample was obtained),
- **D.** subsampling the data by excluding all PJI samples.

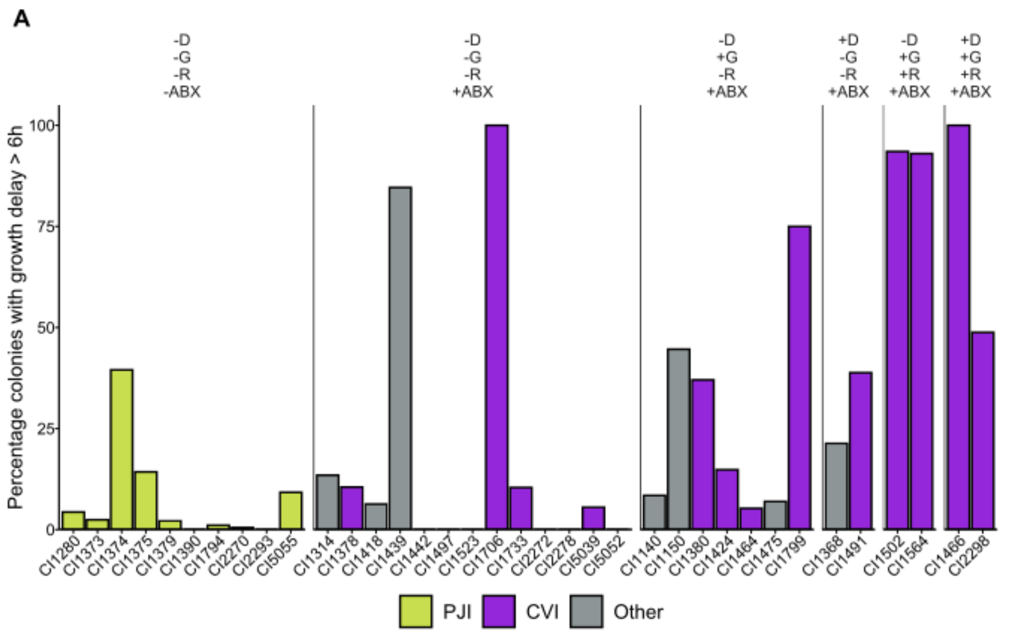

**B**

| Variable | Value | Summary | Univariable p−value | Multivariable p−value |
|---|---|---|---|---|
| Sex | Female | 10 (27.8%) | 0.66 | 0.97 |
| Age (per 10y) | Median (IQR) | 63.0 (37.0−74.0) | 0.29 | 0.23 |
| Immunosuppressed | Yes | 10 (27.8%) | 0.71 | |
| Category | | | 0.021 | 0.51 |
| Reference | PJI | 9 (25.0%) | | |
| compared to | CVI | 15 (41.7%) | | |
| compared to | Other | 12 (33.3%) | | |
| Relapse | Yes | 9 (25.0%) | 0.18 | |
| Bacteremia | Yes | 17 (47.2%) | 0.28 | |
| Abscess | yes | 21 (58.3%) | 0.98 | |
| Implant | Yes | 17 (47.2%) | 0.10 | |
| Betalactam | Yes | 26 (72.2%) | 0.049 | 0.97 |
| GEN | Yes | 11 (30.6%) | 0.004 | 0.48 |
| VAN | Yes | 8 (22.2%) | <0.001 | 0.011 |
| DAP | Yes | 4 (11.1%) | 0.079 | |
| RIF | Yes | 4 (11.1%) | <0.001 | 0.002 |
| Other antibiotics | Yes | 8 (22.2%) | 0.13 | |

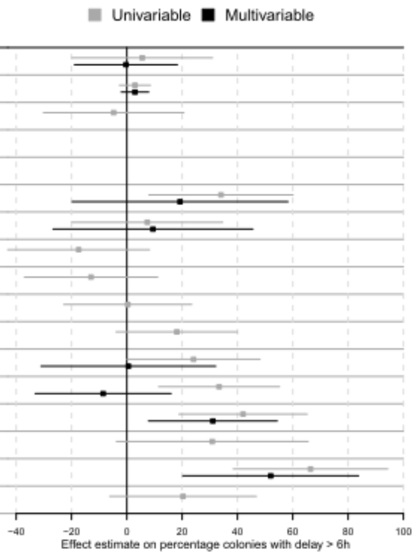

**Suppl. Fig. S7: Multivariable linear regression with alternative outcome A.** Percentage of colonies with a growth-delay larger than six hours from bacterial populations isolated directly from patients. **B.** Association of 14 clinical parameters with percentage of colonies with growth-delay larger than six hours for patient-derived *S. aureus* (n = 36), assessed by univariable and multivariable linear regression. Same as for the Fig. 2 analysis, sex, age, and parameters with a p-value below 0.05 in the univariable model were included in the multivariable analysis. This resulted in a different set of antibiotics to be included in the model: here – instead of GEN, DAP and RIF – Beta-lactams, GEN, VAN and RIF were included. RIF and VAN were associated with an increased proportion (52.0% [20.3, 83.8] and 31.1% [7.84, 54.4] respectively) of colonies with growth-delay larger than six hours. GEN: gentamicin, VAN: vancomycin, DAP: daptomycin, RIF: rifampicin. "Other antibiotics" includes clarithromycin, metronidazole, ciprofloxacin, levofloxacin, tigecycline or tobramycin.

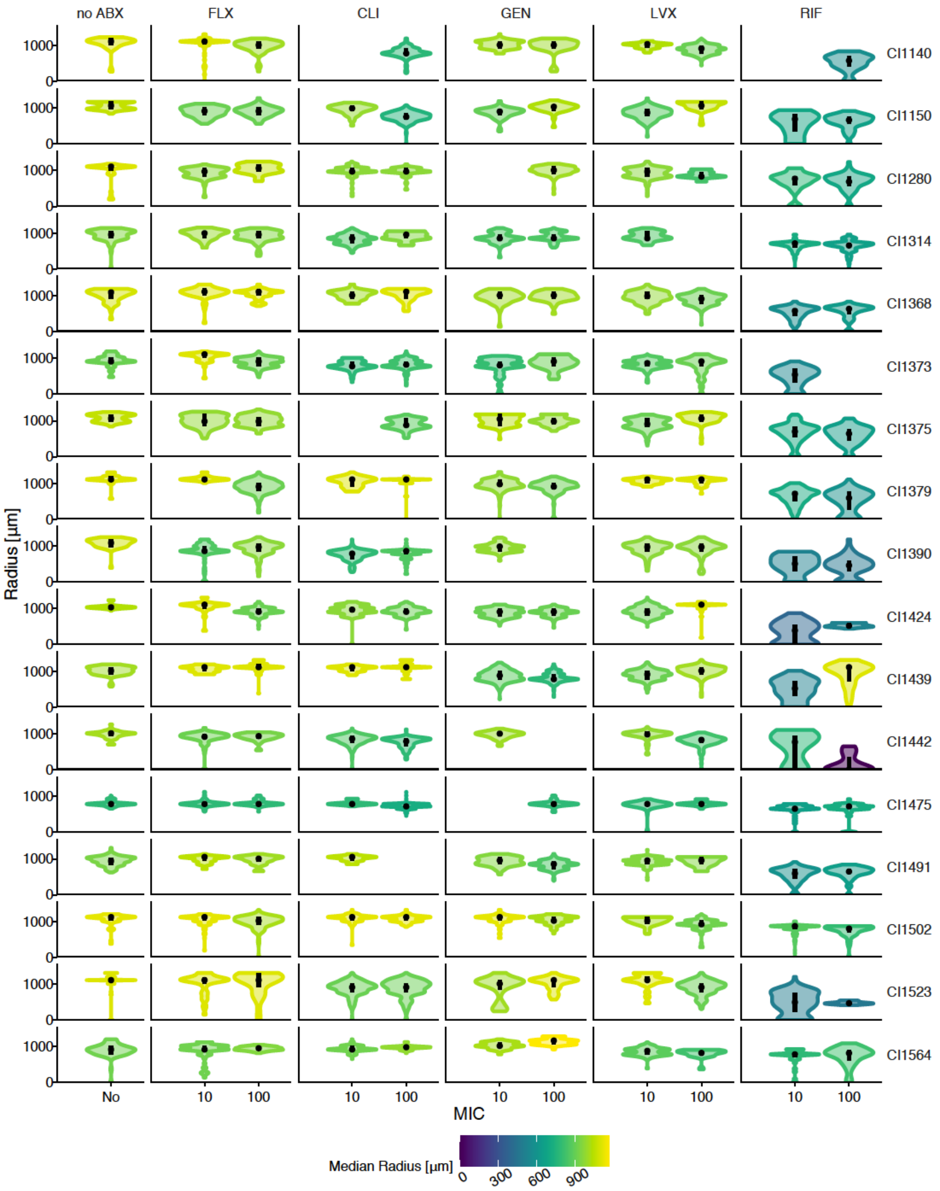

**Suppl. Fig. S8: Colony radius distributions of biofilm derived S. aureus populations**, which had been exposed to five antibiotics (flucloxacillin (FLX), clindamycin (CLI), gentamicin (GEN), levofloxacin (LVX) or rifampicin (RIF)) at two concentrations or a no-antibiotic control (no ABX). The distribution observed on a single plate with minimum 20 and maximum 250 colonies is shown for each treatment. In the few cases in which this colony density criterium was no met, the distribution is not show and was disregarded for downstream analyses. IQR and median are shown in black and color reflects median. These data were used to estimate colony appearance-time distributions, which were then calibrated with the reference appearance-time to obtain growth-delay distributions. Growth-delay distributions for each treatment (all clinical isolates combined) are shown in Fig. 3A.

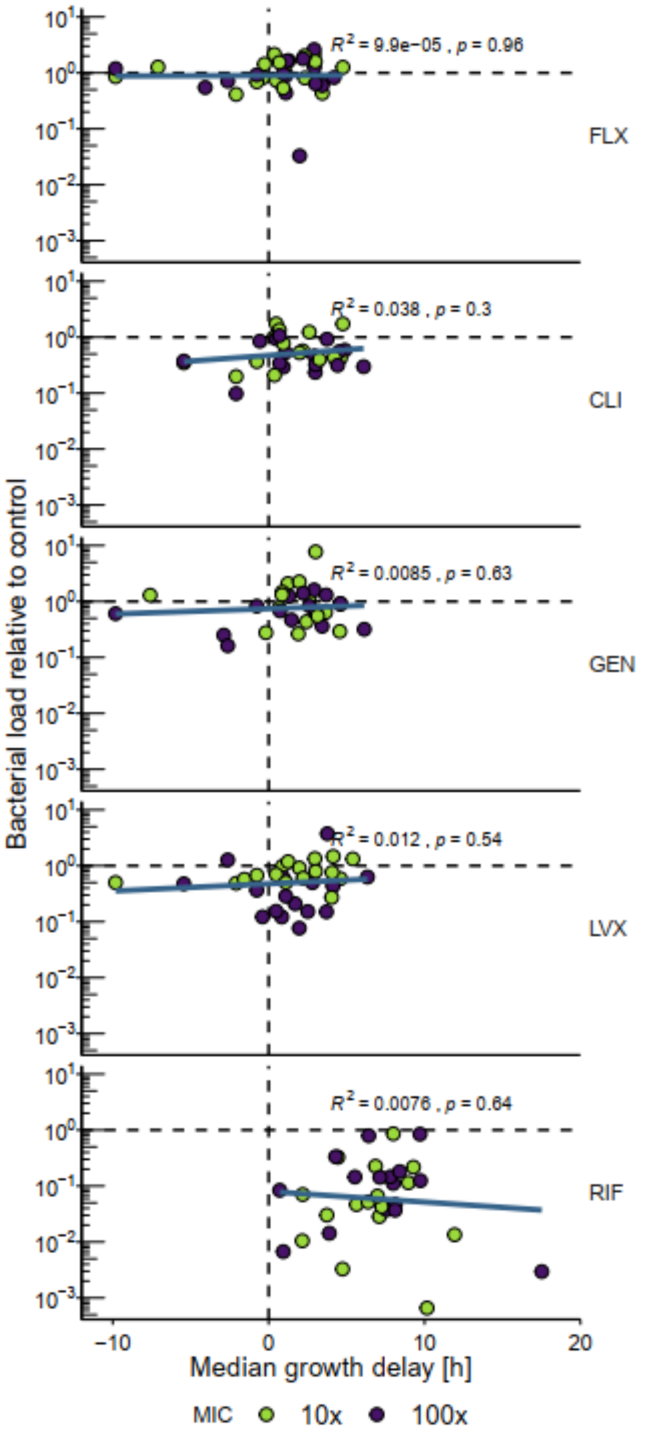

**Suppl. Fig. S9: Relation between median growth-delay and bacterial load** for each treatment (alternative visualization of the data shown in Fig. 3A and B). Each dot represents one clinical isolate. $R^2$ and p-value are based on a Pearson correlation.

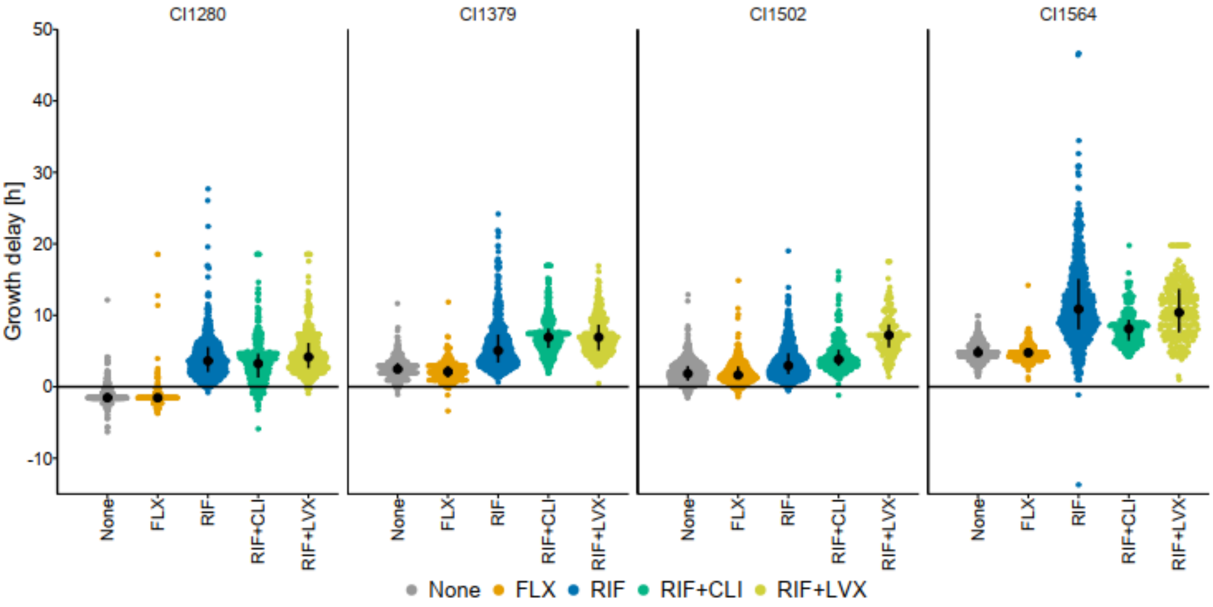

**Suppl. Fig. S10: Growth-delay distributions of biofilm derived S. aureus populations** (alternative visualization of the data shown in Fig. 4B). Each dot represents one colony (data from three biological replicates combined). IQR and median are shown in black.

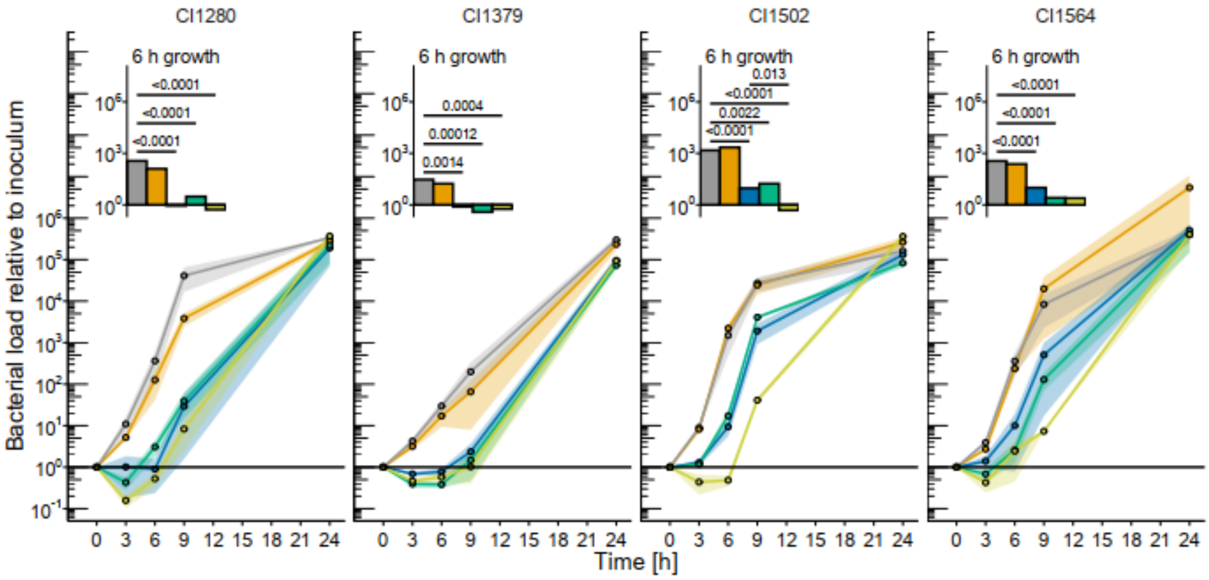

**Suppl. Fig. S11: Regrowth in TSB of biofilm derived S. aureus populations** (same inoculum as in Fig. 4C). Points and shaded area represent mean and standard error of three biological replicates. Relative growth in absence of antibiotics after 6 h was assessed with linear regressions with interaction terms followed by pairwise comparisons computed with estimated marginal means *post-hoc* tests (p-value correction based on multivariate *t*-distribution).

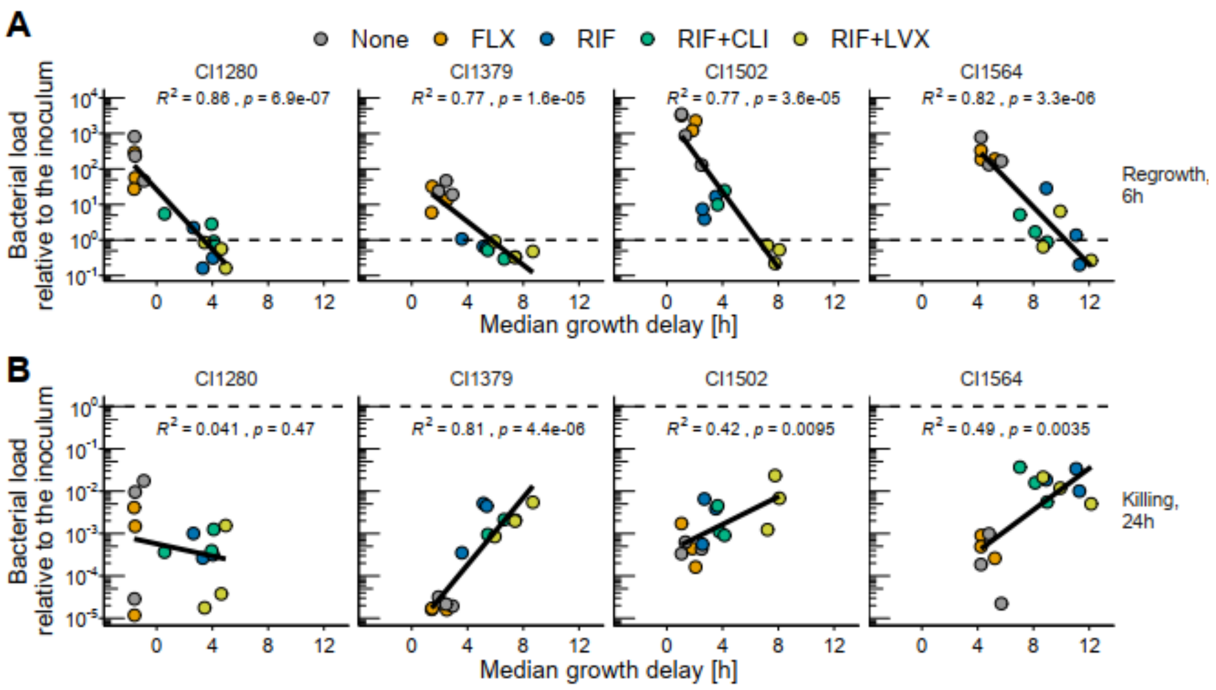

**Suppl. Fig. S12: Relation between median colony growth-delay and bacterial load** (alternative visualization of the data shown in Fig. 4BC and Suppl. Fig S11). Each dot stems from one biological replicate and Pearson correlation per panel is shown.

- **A.** Bacterial load relative to the inoculum after 6 h in TSB without antibiotics
- **B.** Bacterial load relative to the inoculum after 24 h in TSB supplemented with 40x MIC FLX.

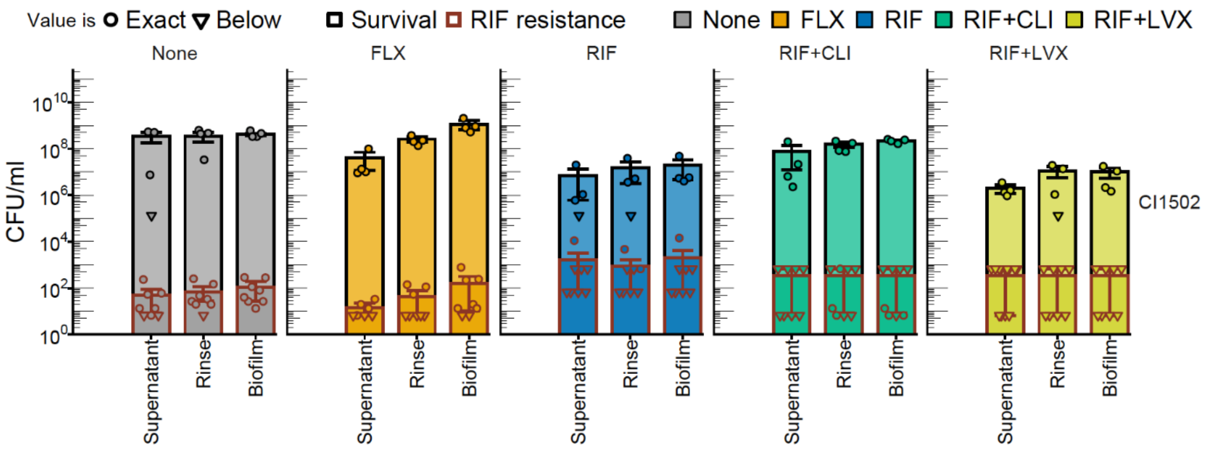

**Suppl. Fig. S13: Bacterial load for each of the three static biofilm fractions** which had been exposed to FLX, RIF or the combination of RIF and CLI or RIF and LVX (black contour) and corresponding load of RIF resistant mutants (overlaid, with a red contour). Mean and standard deviation are shown. Dots represent biological replicates (n = 4, CI1502) and shape indicates if the value was exact or below our detection threshold (Suppl. Method 7).

### Supplementary Tables

**Suppl. Table S1: Minimum inhibitory concentrations** (MIC) of five different antibiotics for each clinical isolate, as determined with ETEST strips (Biomérieux). At the bottom, median MIC, the concentration used in biofilm assays and EUCAST breakpoint values for resistance in *S. aureus* is denoted. If a MIC is above the breakpoint the value is highlighted with bold red text.

| CI ID | GEN [µg/ml] | RIF [µg/ml] | OXA [µg/ml] | LEV [µg/ml] | CLI [µg/ml] |
|---|---|---|---|---|---|
| CI1140 | 0.5 | 0.011 | 0.25 | 0.22 | 0.059 |
| CI1150 | 0.44 | 0.028 | 0.25 | 0.25 | 0.047 |
| CI1280 | 1 | 0.024 | 0.315 | 0.22 | 0.079 |
| CI1314 | 0.44 | 0.01 | 0.625 | 0.188 | 0.016 |
| CI1368 | 0.565 | 0.007 | 0.565 | 0.188 | 0.048 |
| CI1373 | **1.375** | 0.006 | 0.5 | 0.438 | 0.094 |
| CI1374 | 0.25 | 0.008 | 0.25 | 0.125 | 0.064 |
| CI1375 | 0.375 | 0.004 | 0.125 | 0.367 | 0.079 |
| CI1376 | 0.38 | 0.004 | 0.25 | 0.094 | **0.64** |
| CI1378 | 0.25 | 0.006 | 0.38 | 0.125 | **0.64** |
| CI1379 | 0.625 | 0.006 | 0.38 | 0.158 | 0.094 |
| CI1380 | 0.25 | 0.008 | 0.25 | 0.19 | 0.032 |
| CI1390 | 0.22 | 0.007 | 0.875 | 0.158 | 0.086 |
| CI1418 | 0.25 | 0.012 | 0.44 | 0.19 | 0.04 |
| CI1424 | 0.125 | 0.005 | 0.367 | 0.22 | 0.063 |
| CI1439 | 0.22 | 0.005 | 0.44 | 0.533 | 0.063 |
| CI1442 | 0.38 | 0.004 | 0.44 | 0.158 | 0.11 |
| CI1464 | 0.094 | 0.004 | 0.75 | 0.125 | 0.032 |
| CI1466 | 0.25 | 0.004 | 0.188 | 0.094 | **0.625** |
| CI1475 | 0.285 | 0.007 | 0.25 | 0.125 | 0.079 |
| CI1491 | 0.125 | 0.01 | 0.079 | 0.056 | 0.125 |
| CI1497 | 0.25 | 0.012 | 0.25 | 0.19 | 0.064 |
| CI1502 | 0.625 | 0.02 | 0.25 | 0.036 | 0.078 |
| CI1523 | **10** | 0.01 | 1.5 | **1.25** | 0.158 |
| CI1564 | 0.5 | 0.012 | 0.875 | 0.38 | 0.142 |
| CI1706 | 0.38 | 0.008 | **32** | **6** | **>256** |
| CI1733 | 0.5 | 0.003 | 0.5 | 0.25 | 0.125 |
| CI1794 | **64** | 0.006 | **3** | 0.125 | 0.064 |
| CI1799 | 0.25 | 0.006 | 0.19 | 0.19 | 0.094 |
| CI2270 | 0.25 | 0.012 | 0.38 | 0.19 | 0.064 |
| CI2272 | 0.38 | 0.006 | 0.38 | 0.19 | 0.064 |
| CI2278 | 0.25 | 0.006 | 0.19 | 0.19 | 0.064 |
| CI2298 | 0.38 | 0.012 | 0.5 | 0.19 | 0.032 |
| CI2293 | 0.38 | 0.008 | 0.38 | 0.125 | 0.064 |
| CI5039 | 0.25 | 0.008 | 0.25 | 0.125 | 0.047 |
| CI5052 | 0.25 | 0.004 | 0.19 | 0.125 | 0.064 |
| CI5055 | 0.38 | 0.012 | 0.38 | 0.25 | 0.032 |
| **MEDIAN** | **0.38** | **0.007** | **0.38** | **0.19** | **0.064** |
| **USED MIC** | **0.5** | **0.01** | **0.5** | **0.25** | **0.1** |
| **100x MIC** | **50** | **1** | **50** | **25** | **10** |
| **Breakpoint (EUCAST v11.0)** | **1** | **0.5** | **2** | **1** | **0.5** |

**Suppl. Table S2 Characteristics of the sampled infections**. CI: clinical isolate identifier. Patient ID: patient identifier. Category: clinical category of the infection (PJI, Prosthetic joint infection; CVI, cardiovascular infection). Disease: diagnosed disease which was the reason for surgical procedure or other sampling procedure. Sample source: material from which the bacteria were isolated (BAL, bronchoalveolar lavage). Samples were obtained from a surgical procedure in 80% (25/31). Of these, 84% (21/25) were emergency procedures (recently diagnosed infections requiring immediate source control for bacterial load reduction and prevention of organ failure). Only 12% (3/25) were elective procedures and one was performed for a reason other than the ongoing infection. Based on inflammatory laboratory values and radiology, 90% (28/31) patients had an acute inflammatory status at the time of sampling and the remaining three a subacute inflammatory status. CRP (C-reactive protein) and Lc (Leucocyte count): closest available measurement to sampling timepoint. CRP (C-reactive protein) and Lc (Leucocyte count): closest available measurement to sampling timepoint. Time since symptoms: days since symptom onset. >365 indicate a chronic sinusitis case with multiple years of recurring symptoms. Fatal outcome: "yes" indicates death of the patient due to the infection.

| CI | Patient ID | Category | Disease | Sample source | Surgical procedure | Inflammatory status | CRP [mg/l] | Lc [G/l] | Time since symptoms [d] | Fatal outcome |
|---|---|---|---|---|---|---|---|---|---|---|
| CI1140 | ID90 | Other | Spinal abscess | Pus | Emergency | Acute | 283 | 20.3 | 7 | yes |
| CI1150 | ID90 | Other | Spinal abscess | Pus | Emergency | Acute | 308 | 16 | 10 | yes |
| CI1280 | ID87 | PJI | Prosthetic joint infection | Sonicate | Emergency | Acute | 238 | 10.76 | 476 | no |
| CI1314 | ID81 | Other | Septic shock due to pleural empyema | Tissue | Emergency | Acute | 169 | 40.3 | 6 | no |
| CI1368 | ID81 | Other | Septic shock due to pleural empyema | Pus | Emergency | Acute | 43 | 21.5 | 11 | no |
| CI1373 | ID79 | PJI | Septic gonarthritis | Pus | Emergency | Acute | 508 | 12.48 | 1 | yes |
| CI1374 | ID75 | PJI | Prosthetic joint infection | Sonicate | Emergency | Acute | 33.8 | 14.3 | 4 | no |
| CI1375 | ID74 | PJI | Prosthetic joint infection | Sonicate | Emergency | Acute | 222 | 10.94 | 4 | no |
| CI1378 | ID78 | CVI | Native valve endocarditis | Tissue | Emergency | Acute | 96 | 21.3 | 10 | no |
| CI1379 | ID74 | PJI | Prosthetic joint infection | Punctate | Emergency | Acute | 222 | 10.94 | 4 | no |
| CI1380 | ID77 | CVI | Native valve endocarditis | Tissue | Emergency | Acute | 50 | 16.1 | 12 | no |
| CI1390 | ID57 | Other | Soft tissue infection | Tissue | Emergency | Acute | 4.4 | 15.6 | 3 | no |
| CI1418 | ID62 | Other | Folliculitis | Pus | None | Acute | NA | NA | 4 | no |
| CI1424 | ID63 | CVI | Septic pulmonary emboly due to NVE | BAL | Emergency | Acute | 125 | 13.48 | 15 | yes |
| CI1439 | ID58 | Other | Septic shock due to bilateral pneumoniae | BAL | Other Reason | Acute | 245 | 46.8 | 7 | no |
| CI1442 | ID57 | Other | Pneumonia | BAL | None | Acute | 182 | 14.4 | 5 | no |
| CI1464 | ID53 | CVI | Native valve endocarditis | Blood | None | Acute | 147 | 7.66 | 7 | no |

| | | | | | | | | | | |
|---|---|---|---|---|---|---|---|---|---|---|
| CI1466 | ID54 | CVI | PVE, cardiac device infection | Tissue | Emergency | Acute | 16 | 23.97 | 16 | yes |
| CI1475 | ID50 | Other | Meningitis/ventriculits due to pansinusitis | Blood | Emergency | Acute | 47 | 14.35 | 8 | no |
| CI1491 | ID44 | CVI | Native valve endocarditis | Tissue | Emergency | Acute | 26 | 16.22 | 12 | no |
| CI1497 | ID39 | CVI | Suspected cardiac device infection | Sonicate | Elective | Acute | 50 | 10.9 | 22 | no |
| CI1502 | ID36 | CVI | Graft infection | Pus | Emergency | Acute | 55 | 30.48 | 9 | no |
| CI1523 | ID1 | Other | Chronic sinusitis | Nasal swab | None | Subacute | NA | NA | >365 | no |
| CI1564 | ID27 | CVI | Prosthetic valve endocarditis | Sonicate | Emergency | Acute | 106 | 40.76 | 5 | yes |
| CI1706 | ID25 | CVI | Native valve endocarditis | Tissue | Emergency | Acute | 50 | 13.08 | 14 | yes |
| CI1733 | ID21 | CVI | Endocarditis, septic emboli | Pus | Emergency | Acute | 69 | 23.27 | 6 | yes |
| CI1794 | ID9 | PJI | Prosthetic joint infection | Sonicate | Emergency | Acute | 174 | 10.56 | 5 | no |
| CI1799 | ID6 | CVI | NVE, graft and cardiac device infection | Sonicate | Emergency | Acute | 18 | 15.4 | 15 | yes |
| CI2270 | ID1 | Other | Chronic sinusitis | Pus | None | Acute | NA | NA | >365 | no |
| CI2272 | ID1 | Other | Chronic sinusitis | Pus | None | Subacute | NA | NA | >365 | no |
| CI2278 | ID95 | CVI | Cardiac device infection | Sonicate | Emergency | Acute | 52 | 8.6 | 13 | no |
| CI2293 | ID104 | PJI | Prosthetic joint infection | Sonicate | Emergency | Acute | 477 | 12.6 | 5 | no |
| CI2298 | ID105 | CVI | prosthetic valve endocarditis, graft and cardiac device infection | Sonicate | Elective | Acute | 19 | 4.95 | 23 | no |
| CI5039 | ID123 | CVI | Native valve endocarditis | Tissue | Emergency | Acute | 149 | 14.6 | 8 | yes |
| CI5052 | ID159 | PJI | Prosthetic joint infection | Sonicate | Emergency | Acute | 374.8 | 9.5 | 5 | no |
| CI5055 | ID161 | PJI | Prosthetic joint infection | Sonicate | Elective | Subacute | 6.6 | 5.12 | 168 | no |

**Suppl. Table S3: Explanatory variables used in linear regression analyses**. The first 14 variables were included in the univariable and considered for inclusion in the multivariable linear regression analysis (Fig. 2, Suppl. Fig. S6) and two additional technical variables were included in a sensitivity analysis (Suppl. Fig. S7B).

| | Variable | Description |
|---|---|---|
| 1 | Sex | Sex of the patient. |
| 2 | Age | Age of the patient. Note, for better readability of the confidence interval, age divided by ten was used for model calculations. |
| 3 | Immunosuppressed | Indicating if the patient was under any kind of immunosuppressive treatment or suffered from an immunodeficiency disorder. |
| 4 | Category | Clinical category of infection. Either prosthetic joint infections (PJI), cardiovascular infections (CVI) or other infections. “Other” includes variety of infections ranging from soft tissue and skin infections to sinusitis. |
| 5 | Relapse | Indicating if the sample was derived from an infection which was considered a relapse from a previous infection. |
| 6 | Bacteremia | Indicating if the patient had a bacteremia at the time of sampling or shortly before. |
| 7 | Abscess | Indicating if the infection was related to an abscess. |
| 8 | Implant | Indicating if the infection was related to an artificial implant. |
| 9 | Beta-lactam | Indicating if the patient was treated with any kind of beta-lactam antibiotic in the course of the current infection. |
| 10 | GEN | Indicating if the patient was treated with gentamicin in the course of the current infection. |
| 11 | VAN | Indicating if the patient was treated with vancomycin in the course of the current infection. |
| 12 | DAP | Indicating if the patient was treated with daptomycin in the course of the current infection. |
| 13 | RIF | Indicating if the patient was treated with rifampicin in the course of the current infection. |
| 14 | Other antibiotics | Any other antibiotic that was given to < 4 patients of our cohort. Includes: Clarithromycin, Metronidazole, Ciprofloxacin, Levofloxacin, Tigecycline and Tobramycin. |
| 15 | Imaging method | Imaging method used for colony appearance-time determination. Either time-lapse (TL) or endpoint (EP) imaging. |
| 16 | Preparation delay | Preparation delay in days from sampling to bacteria isolation (ranging from 0 to 2). |

**Suppl. Table S3: Clinical data for each sample** considered in regression analyses. The first two column correspond to the clinical isolate and patient identifiers. The following 9 columns correspond to the variables described in Suppl. Table 3. Treatment was subdivided into 6 variables for the model, see Suppl. Table S3. The following two columns correspond to the technical variables described in in Suppl. Table S3. Finally, the last three columns correspond to distinct summary values of the growth-delay distributions. The median was chosen as outcome for the model shown in Fig. 2. Results obtained using "percentage of colonies with growth delay> 6h" as alternative outcome are shown in Suppl. Fig S7B. PJI: prosthetic joint infection; CVI: cardiovascular infection; BET: any kind of beta-lactam antibiotic; GEN: gentamicin; VAN: vancomycin; LEV: levofloxacin; DAP: daptomycin; MET: metronidazole; CLA: clarithromycin; CLI: clindamycin; RIF: rifampicin; TIG: tigecycline; TOB: tobramycin; TL: time-lapse; EP: endpoint; IQR: interquartile range.

| Clinical isolate ID | Patient ID | Sex | Age [y] | Immunosuppressed | Category | Relapse | Bacteremia | Abscess | Implant | Treatment | Type | Preparation delay [d] | Median growth-delay [h] | IQR growth-delay [h] | % colonies with growth-delay >6h |
|---|---|---|---|---|---|---|---|---|---|---|---|---|---|---|---|
| CI1140 | ID90 | m | 65 | no | Other | no | yes | yes | no | BET, GEN, VAN, CLI | TL | 0 | 1.71 | 1.71 | 8.47 |
| CI1150 | ID90 | m | 65 | no | Other | no | yes | yes | no | BET, GEN, VAN, LEV, CLI | TL | 0 | 5.86 | 3.19 | 44.62 |
| CI1280 | ID87 | m | 77 | yes | PJI | no | yes | no | yes | None | TL | 1 | 0.16 | 1.75 | 4.35 |
| CI1314 | ID81 | m | 58 | yes | Other | no | no | yes | no | BET, MET | TL | 0 | -1.04 | 3.35 | 13.6 |
| CI1368 | ID81 | m | 58 | yes | Other | no | no | yes | no | BET, DAP, MET | TL | 0 | 3.12 | 3.09 | 21.36 |
| CI1373 | ID79 | m | 87 | no | PJI | no | yes | yes | yes | None | TL | 0 | 0.48 | 1.94 | 2.45 |
| CI1374 | ID75 | f | 77 | no | PJI | yes | no | no | yes | None | EP | 2 | 5.35 | 5.04 | 39.53 |
| CI1375 | ID74 | f | 74 | no | PJI | no | yes | yes | yes | None | TL | 0 | 0.66 | 1.85 | 14.29 |
| CI1378 | ID78 | f | 29 | no | CVI | no | no | yes | no | BET | EP | 0 | 4.4 | 2.49 | 10.53 |
| CI1379 | ID74 | f | 74 | no | PJI | no | yes | yes | yes | None | TL | 1 | -0.04 | 0.76 | 2.22 |
| CI1380 | ID77 | f | 63 | no | CVI | no | no | no | no | BET, GEN | EP | 0 | 4.99 | 5.48 | 37.04 |
| CI1390 | ID57 | m | 21 | yes | Other | no |  | yes | no | None | TL | 0 | 1.37 | 1.22 | 0 |
| CI1418 | ID62 | m | 20 | yes | Other | yes |  | yes | no | BET | TL | 0 | 2.05 | 1.34 | 6.32 |
| CI1424 | ID63 | m | 36 | no | CVI | no | yes | yes | no | BET, GEN | TL | 0 | 1.7 | 4.18 | 14.81 |

| | | | | | | | | | | | | | | | |
|---|---|---|---|---|---|---|---|---|---|---|---|---|---|---|---|
| CI1439 | ID58 | m | 55 | no | Other | no | no | no | no | BET, VAN, LEV, CLA | TL | 0 | 9.64 | 9.19 | 84.62 |
| CI1442 | ID57 | m | 22 | yes | Other | no | no | no | no | BET | TL | 0 | 0.57 | 1.69 | 0 |
| CI1464 | ID53 | m | 39 | no | CVI | no | | no | no | BET, GEN, VAN | EP | 0 | -2.44 | 4.41 | 5.26 |
| CI1466 | ID54 | m | 60 | no | CVI | no | yes | yes | yes | BET, GEN, DAP, RIF | EP | 0 | 27.16 | 14.3 | 100 |
| CI1475 | ID50 | m | 80 | no | Other | no | yes | yes | no | BET, GEN | TL | 0 | -0.21 | 1.44 | 6.98 |
| CI1491 | ID44 | m | 29 | no | CVI | no | yes | no | no | BET, DAP | TL | 0 | 4.44 | 4.62 | 38.82 |
| CI1497 | ID39 | m | 76 | yes | CVI | no | no | no | yes | BET | EP | 0 | 1.94 | 1.3 | 0 |
| CI1502 | ID36 | m | 73 | no | CVI | no | no | yes | yes | BET, GEN, RIF | TL | 0 | 14.95 | 11.1 | 93.53 |
| CI1523 | ID1 | f | 36 | no | Other | yes | no | no | no | BET | TL | 0 | -0.14 | 0.47 | 0 |
| CI1564 | ID27 | f | 68 | yes | CVI | no | yes | yes | yes | BET, GEN, VAN, RIF, TIG | TL | 0 | 20.22 | 18 | 93.01 |
| CI1706 | ID25 | m | 87 | no | CVI | no | no | no | yes | BET, VAN | EP | 0 | 9.17 | 1.29 | 100 |
| CI1733 | ID21 | f | 71 | no | CVI | no | yes | yes | no | BET, TOB | TL | 1 | 0.6 | 2.5 | 10.42 |
| CI1794 | ID9 | m | 74 | no | PJI | yes | yes | no | yes | None | EP | 1 | 0.51 | 3.12 | 1.1 |
| CI1799 | ID6 | m | 39 | yes | CVI | no | no | yes | yes | BET, GEN, VAN | EP | 0 | 7.89 | 4.92 | 75 |
| CI2270 | ID1 | f | 37 | no | Other | yes | no | yes | no | None | TL | 0 | 2.39 | 0.67 | 0.51 |
| CI2272 | ID1 | f | 37 | no | Other | yes | no | yes | no | BET | TL | 0 | -0.74 | 0.79 | 0 |
| CI2278 | ID95 | m | 80 | no | CVI | no | yes | no | yes | BET | TL | 0 | -1.88 | 0.81 | 0 |
| CI2293 | ID104 | m | 63 | no | PJI | no | yes | yes | yes | None | TL | 1 | -0.24 | 1 | 0 |
| CI2298 | ID105 | m | 52 | no | CVI | yes | no | no | yes | BET, GEN, VAN, DAP, RIF, TOB | TL | 0 | 5.89 | 4.74 | 48.79 |
| CI5039 | ID123 | m | 29 | no | CVI | no | yes | yes | no | BET, GEN | TL | 0 | 1.37 | 1.51 | 5.56 |
| CI5052 | ID159 | m | 73 | yes | PJI | yes | yes | no | yes | BET, GEN | TL | 1 | 0.07 | 0.76 | 0 |
| C5055 | ID161 | m | 63 | no | PJI | yes | no | no | yes | None | TL | 1 | 1.66 | 2.2 | 9.23 |
